# Highly multiplexed molecular and cellular mapping of breast cancer tissue in three dimensions using mass tomography

**DOI:** 10.1101/2020.05.24.113571

**Authors:** Raúl Catena, Alaz Özcan, Laura Kütt, Alex Plüss, IMAXT Consortium, Peter Schraml, Holger Moch, Bernd Bodenmiller

## Abstract

A holistic understanding of tissue and organ structures and their functions requires the detection of molecular constituents in their original three-dimensional (3D) context. Imaging mass cytometry (IMC) makes possible the detection of up to 40 antigens and specific nucleic acids simultaneously using metal-tagged antibodies or nucleic acid probes, respectively, but has so far been restricted to two-dimensional imaging. To enable use of IMC for 3D tissue analyses, we developed mass tomography, which combines quasi deformation-free serial sectioning with novel computational methods. We utilized mass tomography to analyze a breast cancer sample. The resulting 3D representation reveals spatial and cellular heterogeneity, preferential cell-to-cell interactions, detailed tissue-architecture motifs, and the unique microenvironment of a micro-invasion, where micro-metastases clonality is examined, showing that cells arising from the same invasive area, displaying very distinct phenotypes, are all able to produce initial invasive lesions. Mass tomography will provide invaluable insights into the tissue microenvironment, cellular neighborhoods, and tissue organization.

## INTRODUCTION

Tissues and organs are complex ecosystems comprised of numerous cell types arranged in a manner that is inextricably related to function. Understanding tissue functions and pathologies thus requires knowledge of its constituent cells and their states, extracellular matrix proteins, and vasculature in the context of their three-dimensional (3D) architectural arrangement. Historically, tissues have been studied using microscopy modalities, and recently developed methods have enabled various types of 3D tissue analysis (Supplementary Table 1). Confocal 3D microscopy enables subcellular resolution analysis of tissue sections but is limited in the tissue depth that can be analyzed to about 100 μm (*1*). Multi-photon and light-sheet microscopes allow for 3D reconstructions of tissues up to 1 mm tissue depth at lower, yet single-cell resolution (*2, 3*). These 3D microscopy methods have in common the shortcoming that the number of epitopes that can be measured simultaneously is limited, since they rely on fluorescent reports that show high spectral overlap. To enable multiplexed tissue analysis, cycling immunostaining and chromogenic approaches have been implemented. These methods greatly increased the capabilities of fluorescence microscopy to simultaneously detect multiple epitopes and transcripts (*4–7*). The first example of cyclic fluorescent imaging for 3D tissue analysis was published recently (*8*). Since then, other approaches have also been developed, such as the STARmap method, which enables imaging of hundreds transcripts to a depth of 8 μm in 3D (*9*). In addition to fluorescence-based approaches, mass spectrometry-based imaging of epitopes and transcripts is becoming broadly used. In mass spectrometry-based technologies, mass tags, such as a molecule of a defined mass or metal isotopes, are used as reporters on affinity reagents (*10, 11*).

We recently described imaging mass cytometry (IMC), which allows simultaneous detection of up to 40 antigens (*12*) and nucleic acid sequences (*13*) in formalin-fixed paraffin-embedded (FFPE) (*14*), frozen tissue sections (*15*), and in cultured cells (*16*) with subcellular resolution. We also developed the histoCAT and histoCAT++ software toolboxes to enable analysis of cell phenotypes and their interactions in tissues (*14, 17*). To expand IMC-based imaging to the analysis of tissues in 3D, we developed mass tomography (MT). In this method, reported here, the volume and depth of tissue that can be analyzed is only limited by the measurement time.

To demonstrate the utility of mass tomography, we describe the analysis of an invasive ductal breast carcinoma sample. We expanded the histoCAT++ computational toolset for mass tomography, which enabled single-cell, 3D spatially-resolved analysis of these data. The full pipeline, from tissue obtention, to cellular analysis of a complete 3D model can be performed in one week. Using MT we identified a micrometastasis and invasive tumor cells distinguished from surrounding glandular tissue. This micrometastasis was localized to areas of stroma strongly expressing phosphorylated ribosomal protein S6. Such analysis would not be possible in a two-dimensional (2D) setting. This exemplifies how mass tomography applied to breast tumor tissue can shed light on the process of invasion *in situ*. Furthermore, we added to the analysis pipeline machine-learning methods that enabled classification and discovery of cell types and interactions in 3D, which allowed us to identify and map spatially clusters of epithelial basal cells, T cells, and other known phenotypes displaying distinct molecular profiles.

### Generation of 3D models from IMC data

Our MT approach is based on the serial sectioning of a punched tissue cylinder (1mm diameter) from a paraffin block of archival tissue and subsequent 3D reconstruction from 2D IMC data acquired from each individual serial section (Fig. 1A). A major hurdle to achieve 3D tissue reconstruction is the generation of deformation-free sections from paraffin-embedded tissues. We initially assessed different FFPE and OCT tissue-re-embedding and subsequent sectioning methodologies, but major deformations resulted when tissues were sectioned using standard microtomes or cryostats, respectively. Switching to an ultramicrotome with a diamond knife designed for FFPE sectioning overcame the deformation issues during tissue sectioning (*18*). We also found that antigen retrieval at 95 °C can, depending on tissue type, lead to tissue deformation as well. Finally, after testing numerous conditions, we found that antigen retrieval at 80 °C with an incubation time of 80 minutes yielded similar retrieval efficiency for all antigens tested compared to the 40-minute 95 °C protocol while avoiding tissue deformation (Supplementary Fig. 1).

**Figure 1.**
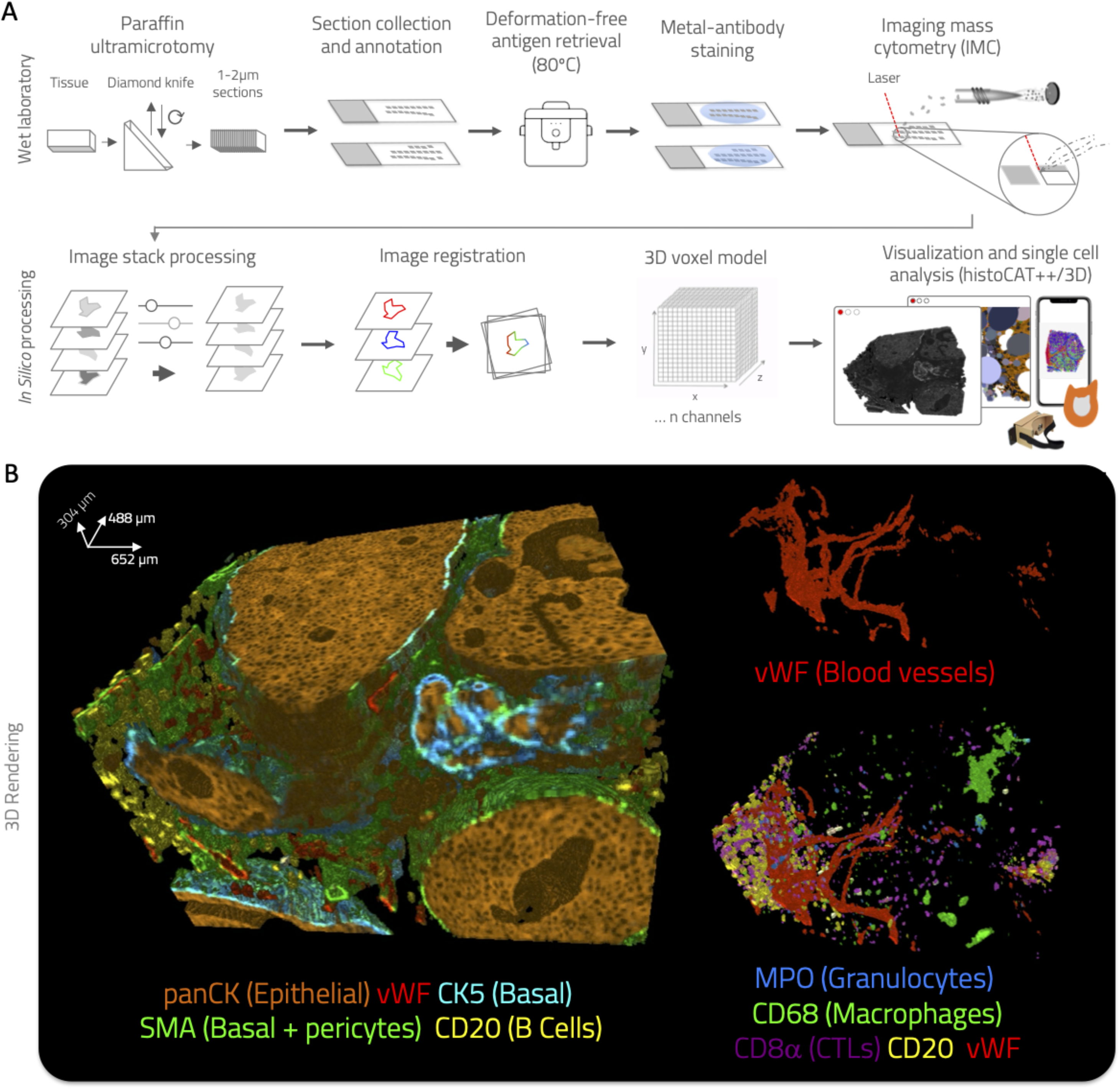
Mass tomography procedure. (**A**) Small rods or blocks of FFPE tissue are cut with a modified diamond-knife using an ultramicrotome into 2-μm sections. Sequential sections are placed on regular microscopy slides. Typically, 40 to 50 sections are placed on each glass slide. After rehydration, tissues are subjected to antigen retrieval, followed by staining with metal-labeled antibodies. All sections are analyzed by IMC. Data are processed computationally to equalize channels, order sections according to the annotation, merge different acquisitions for the same section, and de-noise low intensity channels. Images are then segmented to identify cells, and cells are registered using a novel object-based algorithm. Finally, a full 3D voxel model is assembled and prepared for visualization. (**B**) Examples of renders of different antigens from the same mass tomography 3D voxel model of a breast cancer tumor: (left) pan-cytokeratin (panCK), SMA, vWF, cytokeratin 5 (CK5), and CD20; (upper right) vWF (in red) revealing blood vessels; (lower right) immune cell markers MPO (granulocytes), CD68 (macrophages), CD8α (cytotoxic T cells), CD20 (B cells) in context with the vWF^+^ vasculature.

To illustrate the approach (Fig. 1A, Supplementary Methods), we chose a HER2-positive ductal breast carcinoma sample. After preparation of 156 consecutive slices from the same tissue, we stained the sections with a breast cancer-centric panel of antibodies (Supplementary Table 2) designed to reveal multiple biological aspects of the neoplasm such as tumor cell type, vascularization, immune cell infiltration, cytokeratin composition, proliferation, apoptosis, hypoxia, cell signaling, and collagen deposition. The standard IMC workflow described before was used for imaging of all sections (*12*). Next, to align (i.e., register) the single-section images to perform 3D reconstruction, we implemented a novel alignment method based on single-cell classification. All images were first subjected to single-cell segmentation using a watershed algorithm (Supplementary Fig. 2A). Each cell was assigned a unique identifier, and antibody channel statistics were computed for each cell (Supplementary Figs. 2B-C). We then used a random forest classifier to assign all cells of all sections into one of seven cell phenotypes: luminal epithelial cell, basal epithelial cell, B cell, T cell, macrophage, granulocyte, or stromal cell (Methods, Supplementary Figs. 2D-E). Using these cell labels, we then carried out the registration (Supplementary Methods). Consecutive sections were registered using the cell label match as a metric (Supplementary Fig. 2F). Then, all channels for all sections were rasterized after applying the computed rotations and preprocessing steps to obtain a normalized 3D matrix (Supplementary Figs. 3-4). The retrieved 3D pixel model was visualized and subjected to further processing as necessary for downstream data analysis (Fig. 1B, Supplementary Video 1, Supplementary Fig. 3-5).

The 3D model revealed different constituents of the tumor mass, including the tumor parenchyma (Fig. 1B), where the bulk of the epithelial cells of luminal phenotype reside. These cells express pan-cytokeratin and are negative for the basal markers alpha smooth muscle actin (SMA) and cytokeratin 5 (CK5). The basal layer showed a patchy and discontinuous pattern of the basal markers (Fig. 1B, Supplementary Fig. 6, Supplementary Video 2), indicating that in this particular tumor part of the basal layer has lost structure. Hollow areas within the parenchyma, reminiscent of the original breast gland lumen, are visible within the different carcinoma blobs. The stromal compartment, which contains von Willebrand Factor-positive (vWF^+^) blood vessels, appears heterogeneous in cell composition throughout the volume analyzed (Fig. 1B, Supplementary Video 1). Immune cells, including CD20^+^ B cells, CD8a^+^ cytotoxic T lymphocytes, CD68^+^ macrophages/fibroblasts, and MPO^+^ granulocytes, were detected in this compartment. Microvasculature surrounded by densely packed lymphocytes was visible as were clusters of CD68^+^ macrophages located far from blood vessels and within the hollow areas (Fig. 1B).

### Single-cell data derivation from 3D models

To study the tumor ecosystem of this particular breast cancer sample in depth, we performed 3D single-cell segmentation. We defined the cells within the 3D model using a modified 3D watershed algorithm (Methods, Supplementary Videos 3-4). Each cell was assigned a unique identifier (Fig. 2A, Supplementary Video 4), and statistics for all channels were computed for each 3D cell mask including the antibody signal statistics and morphological data such as volume, shape descriptors, and direct neighbor interactions. These statistics were collated in a cell data catalog with a table format similar to the FCS file format obtained from flow cytometry experiments with cell registries as rows, and channel statistics and coordinates in the 3D model as columns (Fig. 2A).

**Figure 2.**
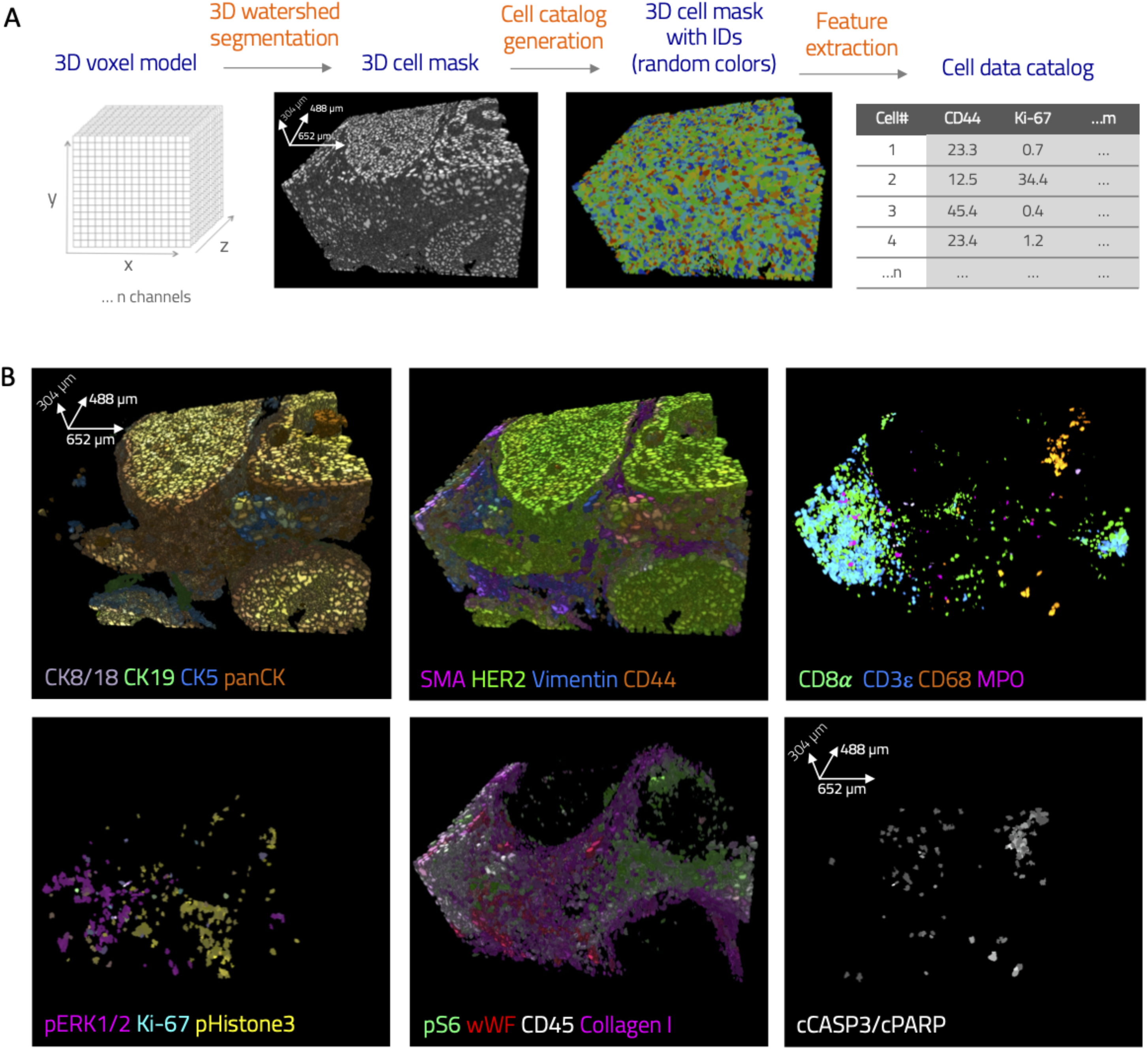
Single-cell analysis of mass tomography data. (**A**) Segmentation and cell catalog generation pipeline. A scalar field (voxel model) is used as an input for a watershed segmentation in 3D. Detected objects are assigned unique cell identifiers. Statistics for all channels and morphological descriptors are calculated for every segmented cell (*N* cells, *M* channels). (**B**) Single-cell protein expression data visualized by rendering over the 3D mask. Multiple markers can be visualized simultaneously with automatically generated or user-defined colors.

To enable the exploration and analysis of the 3D data and cells, we built upon our histoCAT++ analysis platform (*14*) and created histoCAT-3D (Supplementary Fig. 7, Supplementary Video 5). Analysis of our datasets with histoCAT-3D showed that cells that express luminal cytokeratins 8, 18, 19, and cells expressing the basal cytokeratin 5 and other breast cancer epithelial markers such as SMA, HER2, and CD44 delineate the main tumor parenchyma. Different cells expressed these markers in markedly different proportions. This heterogeneity recapitulated our previous findings (*17*). The multiple cell types that make up the stromal compartment were identified by expression of vimentin, collagen I, vWF, and immune markers, including CD45, CD8α, CD3ε, CD68, and MPO. We observed a clear tendency for different types of T cells (e.g., CD45^+^CD3ε^+^CD8α^+^ cytotoxic T lymphocytes and CD45^+^CD3ε^+^CD8α^-^ cells, which are most likely helper T cells) to cluster together around the vWF^+^ microvasculature (Supplementary Videos 1,5). Specific processes of interest were revealed by specific markers, such as Ki-67 and phospho-histone 3, which were often co-expressed and are indicative of proliferation, cleaved caspase-3 and cleaved PARP (cPARP), which are markers of apoptosis, and phospho-ERK1/2 and phospho-S6, which indicate MAPK and PI3K pathway activation, respectively. These observations are in line with conclusions drawn from the raw-voxel data (Fig. 1B). The key advantage of using data post cell segmentation is that the phenotype of each cell in the model can be determined and further cellular downstream analyses can be performed.

### Single-cell data analysis and annotation

In order to comprehensively visualize and annotate the 3D single-cell data we used dimensionality reduction, unsupervised clustering, and supervised classification algorithms in a similar manner to that used for 2D IMC (*14, 17*). These results were added as additional features in columns of the computed cell data table (Fig. 3A). The tSNE embedding clearly separated the main cell types in the sample and showed correspondence with the 3D structure (Fig. 3B-C). Further sub-classifications of cells, using automatic clustering methods, such as FLOCK or k-means, were then mapped directly over a tSNE map and the 3D model (Fig. 3D-E, Supplementary Videos 6-7). To further augment the cell phenotype analysis, we utilized supervised machine learning (SML) to classify the cells. In this breast tumor sample, seven major cell types were defined (basal epithelial cells, luminal epithelial cells, B cells, T cells, macrophages, granulocytes, and other stromal cells), and a random decision tree classifier that learned from all antibody signals was trained by direct interaction with the 3D model (Supplementary Fig. 8, Supplementary Methods) and used to classify more than 50000 cells in the model (Fig. 3F-G). Cell labels from this classification can be overlaid with the unsupervised clustering labels allowing the researcher to perform orthogonal corroboration of phenotypes and clusters identified using unsupervised methods.

**Figure 3.**
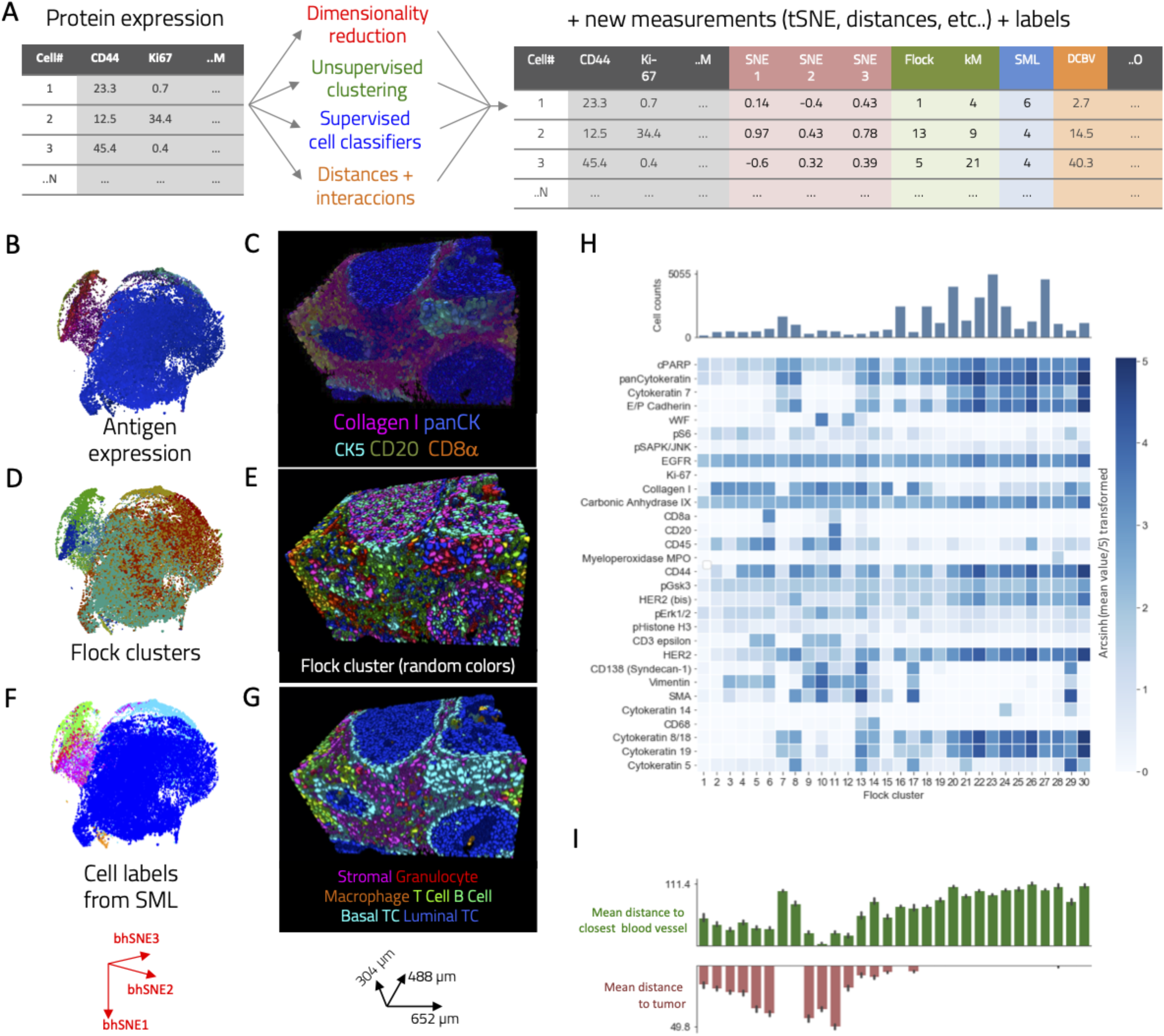
Generation of a comprehensively annotated single-cell 3D atlas. (**A**) Schematic showing the application of multiple methods to augment and annotate the catalog (*N* cells, *M* channels). Here tSNE and principal component analysis were added as new columns (*O* new channels). Cluster identifiers generated using automatic cell clustering methods, such as FLOCK, can be added to the data table as well. Supervised machine learning was used to classify all cells. Topographic information such as distance to particular structures can be calculated by combining the 3D mask with other calculated masks; for instance, the distance to closest blood vessel. (**B-G**) Displays of heterogeneous information types on (B, D, F) 2D projections of a 3D tSNE map and (C, E, G) 3D single-cell maps showing (B, C) color-coded protein expression, (D, E) FLOCK clusters, and (F, G) user-defined cell labels assigned after supervised machine learning classification. For protein expression, color intensity is proportional to protein levels; for categorical data, such as clusters or labels, colors are random and represent cell class. (**H**) Heatmap showing a summary of mean expression of measured markers in discovered cell phenotype clusters after using the FLOCK method. Cluster 11 expression is consistent with B cells (CD20^+^, CD44^+^, CD45^+^) and Cluster 6 with cytotoxic T cells (CD8^+^, CD3^+^); Clusters 5, 9, and 10 likely represent other T cell types. Multiple clusters consistent with tumor cells were identified, underscoring a high heterogeneity of protein expression in tumor cells. Clusters 13 and 14 are characterized by CD68 expression and are likely macrophages. (**I**) Average distance to tumor mask (bottom) or to a blood vessel mask (top) of all FLOCK clusters.

### Phenotypic analysis of the cellular components of the tumor microenvironment

In order to unravel the cellular composition of the tumor in 3D, we proceeded in a manner analogous to that used for 2D multiparametric imaging (*9, 10, 12, 17*). To identify cell-type clusters in an unbiased manner, without forcing the system to a given k (i.e. number of clusters), we used the FLOCK algorithm, which has been used successfully to cluster single-cell data from multiparametric flow cytometry data (*19, 20*). Using FLOCK, cells were categorized automatically into 30 different phenotypic clusters. Phenotypic and spatial analyses of these groups revealed both expected and novel phenotypes (Fig. 3H-I). Certain clusters correspond to known cell types such as B cells (cluster 11) and endothelial cells (cluster 10). Other known cell compartments of greater complexity were assigned to more than one cluster. For example, T cells were assigned to clusters 5, 6, 9, and 10. Cluster 6 contained cytotoxic T lymphocytes, as it is the only one of the four T cell clusters that displays CD8α expression. Clusters 5, 9, and 10 contain other types of T cells, showing similar marker expression, with the exception of vWF and SMA, which were only present in cluster 10 due to the spill-over caused by the close proximity of these cells to blood vessels (Supplementary Fig. 9).

Two clusters, 13 and 14, are consistent with the macrophage phenotype since cells of both clusters express CD68. Interestingly, these clusters are spatially separated (Supplementary Fig. 10A-B). Cluster 13 cells are located within the stroma of the tumor. In contrast, cluster 14 cells are located in hollow areas within the epithelial packages. The cluster 14 macrophages are associated with cPARP-positive epithelial cells (Supplementary Fig. 10C-D), suggesting that these are areas where cells are undergoing apoptosis. Another interesting example of a finding made using this cell atlas is that there are two different clusters of basal cells, which differ in the HER2 and CD44 levels (Supplementary Figs. 11-12, Supplementary Videos 6-7). These clusters map to patches of basal cells in the 3D model (Supplementary Fig. 10). Understanding these different phenotypes in the basal layer may shed light on the early steps of invasion in ductal carcinomas. Furthermore, analysis of immune cell localization revealed enrichment of different types of lymphocytes in the region of a microvascular scaffold, denoted by presence of vWF^+^ cells, with higher concentrations of cells assigned to FLOCK clusters 9 and 11. On the other hand, clusters 5 and 6 showed appeared enriched in a different stromal area near an epithelial DCIS (Figures 3H-I, Supplementary Fig. 9). All these examples demonstrate that the multi-parametric molecular data enabled identification of clearly related populations that differ in location.

The tumor cell continuum is more complex than those of macrophages or T cells and includes 15 FLOCK clusters. These results indicate that our 3D model represents known biology and provides a detailed molecular subclassification of cancer heterogeneity in 3D (Supplementary Figs. 11-14). Such heterogeneity observed within the tumor compartment is in keep with previous findings from our group in a large scale analyses of large cohorts of breast cancer samples (*12, 21*).

### Cellular and environment relationships can be more accurately determined in a 3D model than a 2D model

We hypothesized that 3D analysis would result in higher accuracy in the measurements of distances and the relationships between cells than does 2D IMC. We confirmed this first by measuring the distances between cells and the nearest blood vessel in 2D sections and reconstructed 3D models (Fig. 4A, Supplementary Video 8). The distances measured in 3D differed from those measured in 2D, being distances were always shorter when measuring in 3D (Fig. 4B). This was expected since in single 2D planes the features above and below the plane cannot be detected.

**Figure 4.**
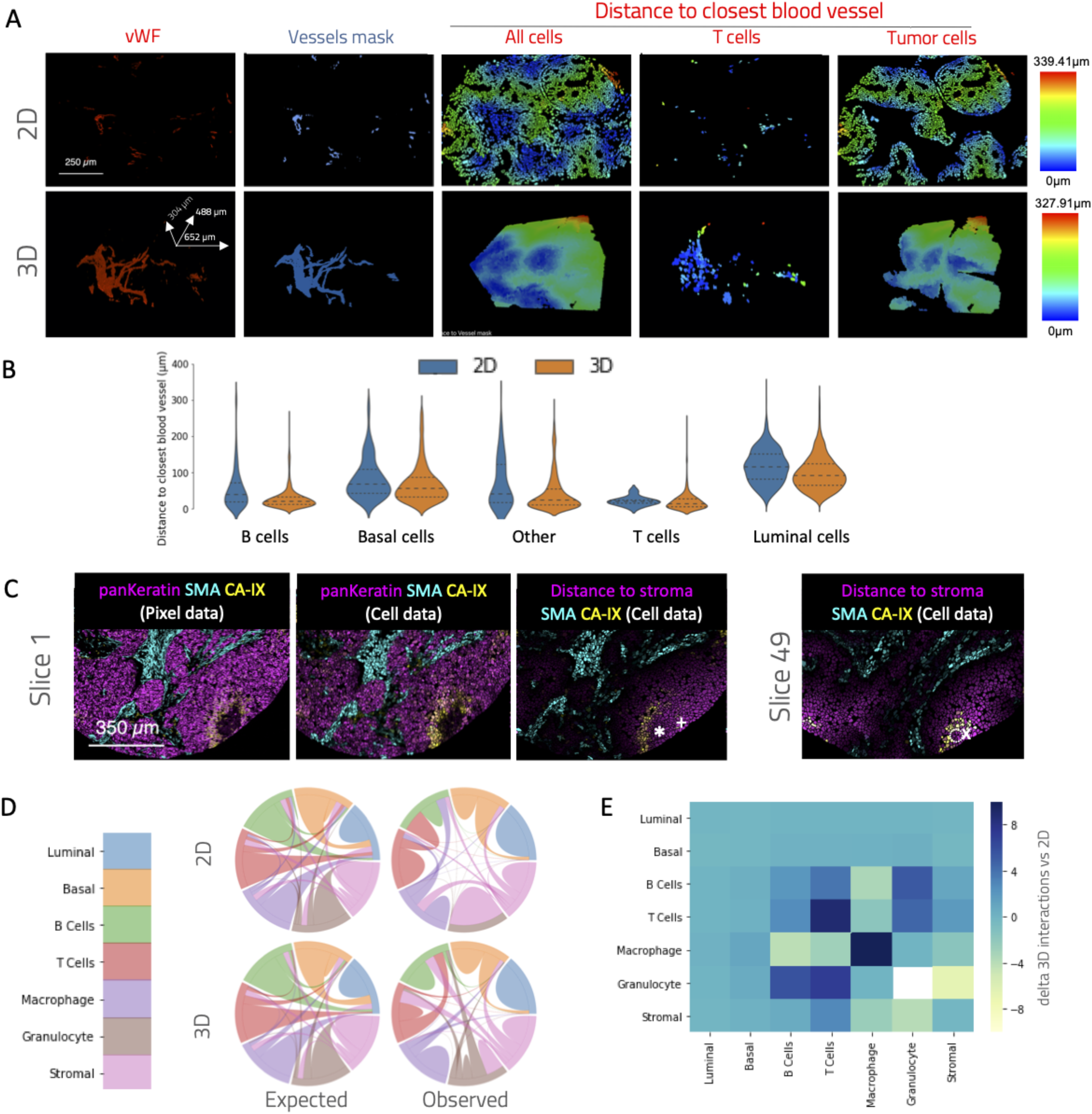
Distance measurements accuracy for 3D versus 2D models. (**A**) Cell distance-to-vessel measurements in 2D (top) and 3D (bottom) models. Left to right the panels show the chosen proxy marker signal, the generated binary masks for vessels, the distance of each cell to the vessel mask in a heat-color scale, and distances to vessel color-coded in heat scales for T cells and tumor cells. (**B**) Quantification of distance from a cell to a vessel in 2D and 3D for selected annotated cell types. (**C**) Slice 1 and slice 49 from a 3D model containing a hypoxic area. Hypoxic centers are marked with asterisks and Xs in slices 1 and 49, respectively, and distance to stroma is marked with + and - for slices 1 and 49, respectively. Markers for hypoxia (carbonic anhydrase IX), stroma (SMA), tumor parenchyma (pan-cytokeratin), and distance are displayed in several color combinations in raw pixel data (leftmost panel) or cell data after segmentation in 2D (all other panels). (**D**) Pairwise cell type interaction chord diagrams (3D vs. 2D and observed vs. expected frequencies). (**E**) Heatmap matrix showing differences in interactions scores between 3D and 2D.

In another tissue sample analyzed, stack images were analyzed one-by-one in 2D as well as in 3D. This tissue displays a hypoxic area in the center of an enlarged epithelial compartment as denoted by positive carbonic anhydrase IX (CAIX) staining. In 2D, a mismatch between the center of the hypoxic area and the farthest point to stroma (oxygen source) at different depths of the tissue was observed (Fig. 4C). In the 3D reconstruction, however, it is apparent that a strip of stromal tissue penetrates into the tumor through half of the depth or the measured volume, explaining the apparent contradiction from 2D (Supplementary Video 9). Thus, the 3D analysis shows that the farthest points from the stroma are the most hypoxic, as expected, whereas this correlation is elusive in the 2D analyses, depending on the depth of the section evaluated.

Finally, we tested whether cell-to-cell interactions are more faithfully represented in 3D than in 2D. To address this we computed a graph of cell interactions and then computed the interaction frequencies amongst all the previously defined cell types for each section in 2D and for the 3D model (Figs. 3 and 4D). Both the expected (predicted number of interactions amongst all cell types if randomly shuffled provided the abundances of each cell type) interactions and the actual, observed interaction rates were calculated for 2D and 3D datasets (Fig. 4D). The differences between the 3D and 2D observed interactions were also quantified (Fig. 4E) (*17*). We could observe that 3D interaction analysis yielded a different cell interaction picture compared to 2D. Examples of this are a greater homotypic interaction amongst macrophages (7.9-fold higher measured in 3D vs. 2D) and a decreased heterotypic B cell to macrophage interaction (5.1-fold lower in 3D vs. 2D). Plausible explanations for these differences are under-sampling in the 2D space and the existence of interactions that occur in constrained directions (e.g., lymphocytes tethered to blood vessels).

### Pseudo-dynamic process analysis

The FFPE tissues analyzed using IMC represent a snapshot in time. Recent work presented approaches to determine a pseudo-time dimension from snapshot single cell data (*22, 23*). Since many dynamic processes in biology follow a spatial-temporal relationship, it should also be possible to infer their chronology from a structural trace. Given that 3D models provide a complete picture of spatial-temporal processes, the data presented here should be ideal for such an analysis.

Indeed, examination of our 3D model indicates a striking example of what can be hypothesized as a spatially detectable dynamic process: Sequential stack reconstruction of one region revealed a micro-invasive lesion and a stream of invasive tumor cells (Fig. 5A, Supplementary Video 10). The distal part of the tumor in the Z axis contained no invading cells, and the tumor cell packages had smooth tumor to stroma boundaries. Toward the center of the Z axis and through the proximal area, invading epithelial cells, emerging from a localized protrusion, were observed (Fig. 5A-B). These invading cells form an inverted cone. As the distance to the point of scape from the tumor increases, invading cells form larger structures that may be incipient micrometastases (Fig. 5C-D). Moreover, we observed that the detected micrometastases in the inverted cone increase in size proportionally to the distance to the protrusion, which we hypothesize is their point of origin (Fig. 5E). This is in agreement with the hypothesis that motile tumor cells escaping tumor structures develop into metastases as they migrate away from the origin (*24*), constituting the incipient lesions that develop into invasive carcinomas. Such correlation requires careful validation with orthogonal methods. Yet, 3D analysis is a valuable for hypotheses generation for such pseudo-dynamic processes, for instance, the aforementioned tumor invasion.

**Figure 5.**
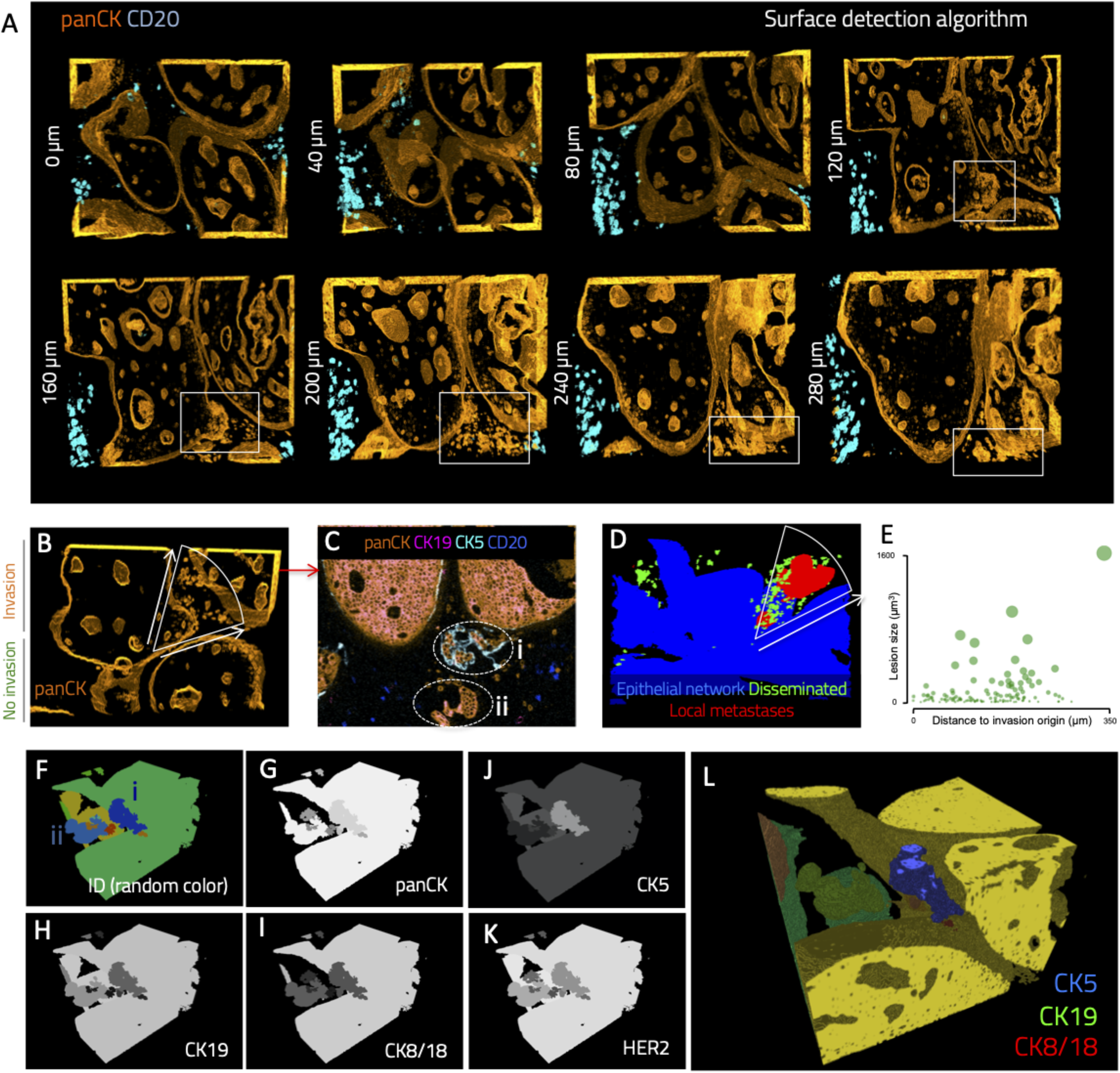
Observation of an invasive process using mass cytometry. (**A**) Sequential tomographic imaging of tumor parenchyma using a surface-detection algorithm. A protrusion in the surface of a DCIS-like structure is visible around 120 μm above the model floor level, and a stream of invading tumor cells are observed between the protrusion and the stroma. A large invasive structure is observed around 240 μm from the floor of the model. (**B**) Projection orthogonal to that of panel A showing the parenchymal protrusion and the area invaded by single tumor cells. (**C**) A 2D projection toward the ceiling of the model shows invasive structures formed at the opposite side of the invading stream. Cytokeratin expression is heterogeneous in different lesions (marked i and ii). (**D**) Epithelial segments are classified as belonging to the tumor network (blue), single disseminated cells (green), or local metastases (red). (**E**) Bubble graph showing pseudotime/distance-to-invasion-origin versus size of invasive lesion (**F**) 3D mapping of local metastases i (blue) and ii (light blue) identified in panel C. (**G-K**) Mean expression of different epithelial markers in the segmented epithelial bodies shows high variability of expression in metastases and the tumor epithelial network. (**L**) Color composite for mean protein expression for CK5, CK19, and CK8/18 in the segmented epithelial bodies.

Due to the multiparametric nature of IMC, it is possible to investigate which tissue components change during the metastatic process. For instance, the invading tumor cells described above seem to migrate toward an area that is enriched in stromal cells that express high levels of phospho-S6 (Supplementary Video 11). It is important to underscore that, in this setting, local micrometastases can be differentiated from sectioned fragments of tumor structures with confidence, as given the 3D information, it is possible to determine whether a given epithelial body is disconnected from the tumor network or not (Fig. 5D-E); this is impossible in 2D cuts (Fig. 5C). Strikingly, two adjacent metastases (Fig. 5F, *i* and *ii*) showed very different levels of numerous molecules (Fig. 5G-L), most notably cytokeratins. Both lesions express low levels of epithelial markers CK8/19 and HER2. Yet, one lesion displayed a higher level of expression of the basal marker CK5 and a lower level of expression of CK19 compared to the other and to the tumor network. These results highlight the suitability of mass tomography, by combining highly parametric molecular analysis with 3D spatial reconstruction, to further our understanding of complex dynamic processes such as of the clonal evolution of cells during the processes of invasion and metastasis. In this example, we clearly show that early micro-metastases with the same origin can drastically differ in their cytokeratin expression, supporting a multi-clonal theory of metastasis (*24*).

### Data visualization and analysis

Enhanced data visualization techniques with user-defined semantics are needed to enable interrogation of 3D multiparametric datasets. Thus, we have developed features to efficiently combine all the available information together with additional annotations and metrics. Once collated, the information can be collectively visualized and queried as a cell atlas, using strategies to combine molecular data with labels and voxel, area, or cell masks. Using this approach, higher-level tissue structures, such as blood vessels or collagen beds, can be delineated and annotated as well. Examples of a 3D cell-rendering that combines raw cell data statistics from SMA with a label annotation (Fig. 6A) and voxel maps of vessels and its corresponding binary mask (Fig. 6B, Supplementary Video 8) are shown. Topological relationships can be determined; for example, the distances from every cell to the closest blood vessel can be measured (Fig. 6B). We also implemented different visualization modalities, such voxel-based, isosurfaces (Supplementary Fig. 15, Supplementary Video 12), spheres, and color stripes and patterns (Fig. 6C-D, Supplementary Video 13). The latter is a powerful approach that has been already been applied to microscopy 3D/5D datasets to augment the amount of data that can be simultaneously be visualized in 3D (*25, 26*). Furthermore, we have transferred this 3D cell atlas to a mobile phone application (histoCAT-mobile, Supplementary Video 14) that uses augmented reality. With this app, a researcher can colorize the channels, interact with the cells, and train cell classifiers (Fig. 6E). In the future, these capabilities will enable for an easy and efficient, cloud-based, crowd-sourced data analysis, a modern technique for quality labeling of data for machine learning that leverages the principle of the “wisdom of the crowd”(*27*).

**Figure 6.**
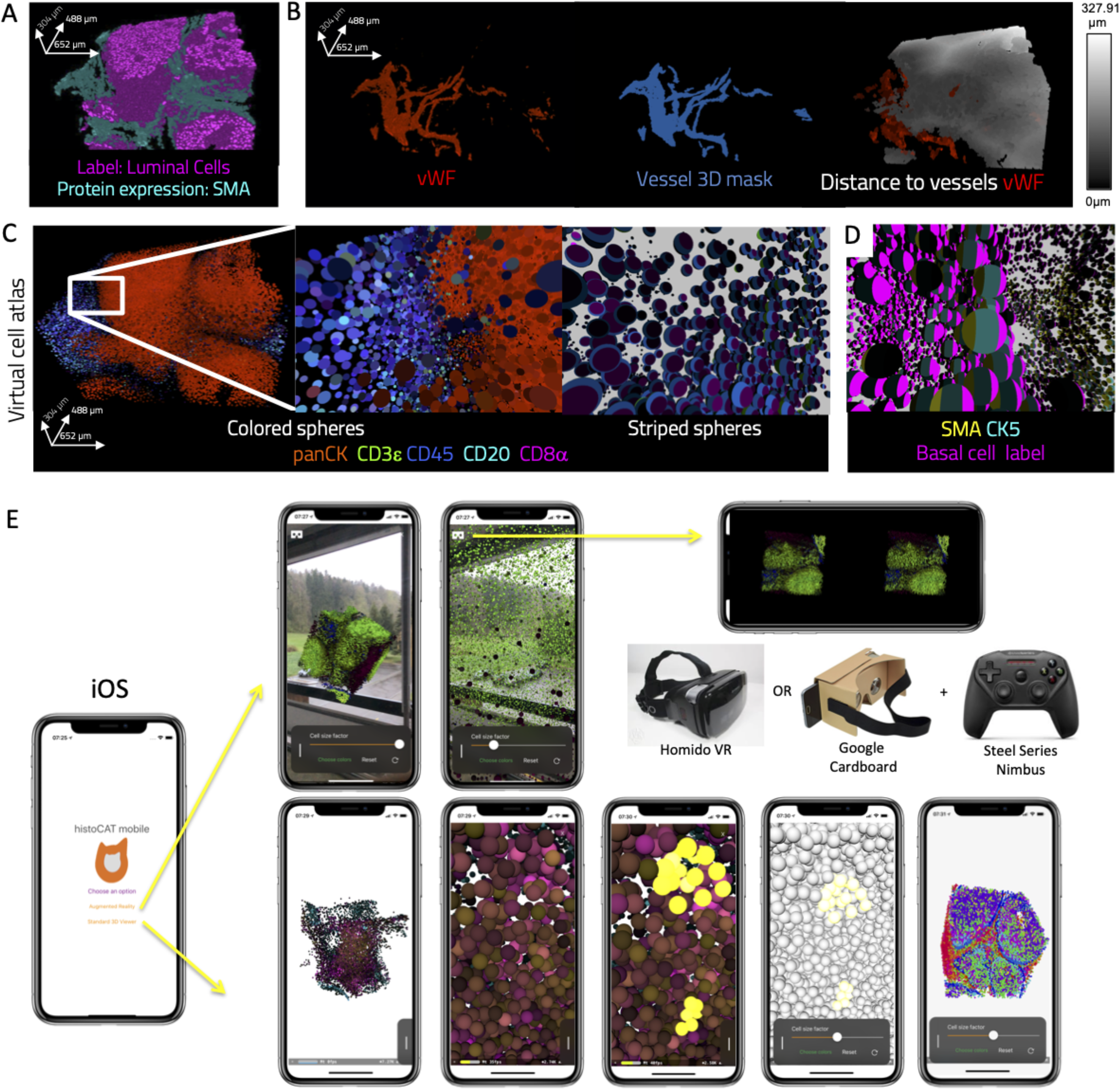
Cell atlas visualization in 3D and augmented reality. (**A**) Simultaneous representation of measured data, here for SMA, and derived data, which in this case is the luminal cell label. (**B**) vWF signal (left) was transformed into a binary mask representing blood vessels (middle). The mask was subsequently used to calculate the distance to closest blood vessel for every cell with measured vWF signal represented simultaneously with the derived distance data with user-defined colors (right). (**C**) Generation of a virtual-shaped atlas. Arbitrary shapes (e.g., spheres), colors, sizes, and patterns (stripes) are used to increase the number of representable variables that can be visually interpreted. (**D**) A sphere model of the tumor with left and center stripes showing SMA and cytokeratin 5 expression, respectively (intensity proportional to protein levels), and right stripe showing basal cell label (full intensity or black for other cell types). (**E**) histoCAT-mobile, an iOS application featuring features augmented reality, virtual reality, and interactive 3D rendering of the cell model. The app allows custom multicolor visualization of channels, and immersive augmented and virtual reality experience to explore and annotate the cells in the tumor. The application is available for download for iPhone 8 or later at: https://itunes.apple.com/us/app/histocat/id1439254241

## CONCLUSION

In summary, we show that multi-parametric tomographic analysis of tissues is possible using mass tomography, with no theoretical size limit. IMC was used to analyze an FFPE archive tissue sample using a serial sectioning method. To analyze the generated data, histoCAT-3D and histoCAT-mobile were developed to allow cell- and tissue-wide segmentation, cell model construction, cell phenotype discovery and annotation, topological data calculation, machine learning-based dataset enrichment, and visualization of the data. The molecular and cellular atlases that can be constructed using MT will enable visualization and analysis of detailed tissue architecture and lineage- and cell-communication mechanisms in native 3D contexts for any tissue type. Some computational and technical aspects still need to be improved. For instance, acquisition time is long (four full days for the presented data), hence, development of hardware with higher ablation speed will allow both shorter acquisition times or alternatively, better resolution, such as 0.5μm in all dimensions (adjusting the section thickness to 0.5μm too). Furthermore, better cell segmentation and channel cross-talk calculation algorithms will reduce the signal spill-over (*28*), or could increase practically the acquisition speed by allowing ablations schemes with lesser resolution, followed by computational processing with tools such as generative adversarial convolutional neural networks (GANs), which can virtually increase the resolution (*29*).

## Supporting information

IMAXT Consortium members

## Supplementary Figure Legends

**Supplementary Fig. 1.** Slice deformation after 95 °C and 80 °C heat-induced antigen retrieval. Top row shows two consecutive slices in white, and the corresponding overlay using red and cyan after a 95 °C, 40-minute antigen retrieval treatment in Tris-HCl buffer. Bottom row shows an equivalent experiment with 80 °C treatment for 80 minutes.

**Supplementary Fig. 2.** Slice registration strategy. (**A**) Each individual IMC image was segmented using a watershed algorithm. (**B**) All channel intensity statistics (features) per cell were calculated using the cell segmentation mask. (**C**) Example cell data for one slice. All features are plotted over the segmentation mask using a white scale for detected protein intensity. (**D**) Interactive cell labeling over the segmentation mask. (**E**) Cell labels obtained using the classifier trained during interactive cell labeling. (**F**) Example of cell label-based registration.

**Supplementary Fig. 3.** 3D voxel model generation pipeline. (**A**) Example overlay of iridium signal from five consecutive slices before registration. (**B**) Example overlay of iridium signal from five consecutive slices after registration. (**C**) Detail of aligned slices with pan-cytokeratin signals in red and cyan on consecutive slices. (**D**) Full 156-slice stack SMA signal before registration. (**E**) SMA signal in full stack after registration. (**F**) Top and side view in 3D of the SMA staining, which labels the basal layers and vessel walls forming a complex mesh. (**G**) A subset of the stack shows correct alignment of the SMA signal. (**H**) Example cube render of a tumor after selection of a region of interest to ensure the 3D volume was generated from an area that contains information from all the slices.

**Supplementary Fig. 4.** Signal from multiple channels after generating the 3D voxel model using an intensity scale.

**Supplementary Fig. 5.** Example of 3D model image processing. Color render of raw voxel data in indicated channels (values obtained by rasterization after applying the rotation transforms are lineally interpolated). Bottom, same render after performing a per channel Gaussian convolution in 3D.

**Supplementary Fig. 6.** Tumor basal layer showing expression of two breast gland basal epithelial markers. SMA expression is uniform within the basal layer, whereas expression of cytokeratin-5 is variable.

**Supplementary Fig. 7.** histoCAT-3D software architecture.

**Supplementary Fig. 8.** Example image of labeling tool for the 3D model. Gray cells are unlabeled, and colored cells have been selected by the researcher and assigned defined phenotypes.

**Supplementary Fig. 9.** Differential immune cell distribution over the tumor stroma. (A) Expression of blood vessel (vWF), fibroblast (collagen I), and immune cell (CD45, CD8a, CD3e, and CD20) markers in the 3D model. (B) Identity color map for clusters 5, 6, 9, and 11. (C) A white ellipse shows cumuli of cells belonging to clusters 5 and 6 in the non-vascularized area of the stroma. (D) A yellow ellipse denotes the vascularized area, marked by vWF+ cells. (E) The same yellow ellipse shows cumuli of cells belonging to the clusters 9 and 11 in this vascularized area.

**Supplementary Fig. 10. Phenotypic analysis of distinct macrophage populations in the 3D model**. (**A**) Clusters 13 and 14 are localized in different topological sites (stromal for cluster 13 and hollow areas within parenchymal tissue for cluster 14) as indicated by (**B**) pan-cytokeratin expression, (**C**) macrophage (CD68) marker expression, and (**D**) cPAPR expression indicative of apoptotic cells.

**Supplementary Fig. 11.** Example of cluster discovery combining supervised and unsupervised machine learning-driven cell labeling. (**A**) 2D projection of a 3D-tSNE map (see also Supplementary Video 5) with cell labels obtained from a trained random forest classifier. A yellow oval indicates an area in the tSNE plot enriched with basal epithelial cells. (**B**) 3D mapping of the cell labels shown in (A). (**C**) Same tSNE projection as in (A) with labels for clusters obtained after using the *k*-means clustering algorithm (*k*=15). The area marked in (A) has two main *k*-means clusters, marked as “cyan cluster” and “blue cluster”. (**D**) 3D mapping over the cell volume of the *k*-means clusters. Arrows indicate examples of the basal localization in DCIS packages for both “cyan” and “blue” *k*-means clusters.

**Supplementary Fig. 12.** Heatmaps showing mean values for all the channels analyzed in all 15 *k*-means clusters and the cells classified into 7 classes using supervised machine learning (SML) obtained from the cell data. Blue bar graphs show the number of events in each bin, green graphs the average distance to the closest blood vessel, and red bars the average distance to the tumor parenchyma.

**Supplementary Fig. 13.** Clustermap showing mean values for all the channels analyzed in all 30 FLOCK clusters obtained from the cell data.

**Supplementary Fig. 14.** Clustermap showing mean values for all the channels analyzed in all 15 *k*-means clusters obtained from the cell data.

**Supplementary Fig. 15.** An example of isosurface rendering of the 3D cell data model.

## SUPPLEMENTARY VIDEOS

**Supplementary Video 1:** Mass tomography rendered voxel data. http://www.bodenmillerlab.org/catena_et_al_mass_tomography/SuppVideo1.mp4

**Supplementary Video 2:** SMA (blue) and cytokeratin 5 (green) markers in voxel 3D reconstruction shows patchy cytokeratin 5 coverage within the epithelial basal membrane. http://www.bodenmillerlab.org/catena_et_al_mass_tomography/SuppVideo2.mp4

**Supplementary Video 3:** Detail of nuclear signal (iridium 191) in 3D. http://www.bodenmillerlab.org/catena_et_al_mass_tomography/SuppVideo3.mp4

**Supplementary Video 4:** Detail of cell mask obtained after performing 3D segmentation over the nuclear signal shown in Supplementary Video 2. http://www.bodenmillerlab.org/catena_et_al_mass_tomography/SuppVideo4.mp4

**Supplementary Video 5:** Cell model showing expression of different measured markers. http://www.bodenmillerlab.org/catena_et_al_mass_tomography/SuppVideo5.mp4

**Supplementary Video 6:** 3D tSNE map showing examples of different measured markers, cluster labels, and labels from a supervised random forests classifier. http://www.bodenmillerlab.org/catena_et_al_mass_tomography/SuppVideo6.mp4

**Supplementary Video 7:** FLOCK, *k*-means, and supervised classification labels displayed over the 3D tumor cell model. http://www.bodenmillerlab.org/catena_et_al_mass_tomography/SuppVideo7.mp4

**Supplementary Video 8:** Blood vessel structural mask and resulting computation of distance to vessel mask for all cells or different cell subsets. http://www.bodenmillerlab.org/catena_et_al_mass_tomography/SuppVideo8.mp4

**Supplementary Video 9:** Section of a hypoxic area of a tumor showing distance to stromal areas, a stromal marker (SMA), and a hypoxia marker (carbonic anhydrase IX). http://www.bodenmillerlab.org/catena_et_al_mass_tomography/SuppVideo9.mp4

**Supplementary Video 10:** 3D axial tour through a 3D tumor model displaying epithelial and B lymphocyte surfaces. In the middle of the video, from the center and toward the lower right quadrant, a protrusion from which disseminated cells spring is observed. http://www.bodenmillerlab.org/catena_et_al_mass_tomography/SuppVideo10.mp4

**Supplementary Video 11:** 3D axial tour through a 3D tumor model displaying epithelial and B lymphocyte surfaces and pS6 signal. http://www.bodenmillerlab.org/catena_et_al_mass_tomography/SuppVideo11.mp4

**Supplementary Video 12.** Isosurface rendering of the 3D cell data model. http://www.bodenmillerlab.org/catena_et_al_mass_tomography/SuppVideo12.mp4

**Supplementary Video 13:** 3D tumor cell atlas showing spherical representation of cells with blended or striped marker colorings. http://www.bodenmillerlab.org/catena_et_al_mass_tomography/SuppVideo13.mp4

**Supplementary Video 14.** histoCAT-mobile demo. http://www.bodenmillerlab.org/catena_et_al_mass_tomography/SuppVideo14.mp4

## ACKNOWLEDGEMENTS

We would like to thank Stefanie Engler and Andrea Jacobs (University of Zürich) and Susanne Dettwiler and Fabiola Prutek (Tissue biobank, University Hospital Zürich) for technical support. Thanks to Professor Roger Albert Wepf (University of Queensland, Brisbane), Helmut Gnaegi (Diatome AG, Nidau, Switzerland), and Urs Ziegler, Andres Käch, and Gery Barmettler from the University of Zürich for their invaluable guidance with the paraffin ultramicrotomy procedure. Special thanks too to Jonas Windhager and Vito Zanotelli for their extraordinary advice on computational analysis. This work was supported by the Swiss National Science Foundation (SNSF) R’Equip grant 316030-139220, an SNSF Assistant Professorship grant PP00P3-144874, the PhosphonetPPM and MetastasiX SystemsX grants, and funding from the European Research Council (ERC) under the European Union’s Seventh Framework Programme (FP/2007-2013)/ERC Grant Agreement no. 336921.

## AUTHOR CONTRIBUTIONS

RC and BB conceived the study. RC designed mass tomography, performed all experiments, analyzed the data, and developed histoCAT-3D and histoCAT mobile software packages. AP, LK, and AO assisted with experimentation. PS and HM contributed clinical samples and scientific guidance. RC and BB wrote the manuscript. All authors have read and edited the manuscript.

