## Supplementary Figures for "Highly multiplexed molecular and cellular mapping of breast cancer tissue in three dimensions using mass tomography"

| | Max number of antigens/targets | Resolution | Maximum depth/thickness | References |
| --- | --- | --- | --- | --- |
| Confocal microscopy | ~8 | nanometric/submicrometric | ~100 µm | 1 |
| Two-photon microscopy | 3/4 | submicrometric | ~1000 µm | 2 |
| Electron microscopy tomography / Array tomography | 2 | nanometric | Unlimited. Few micra in practice | 3 |
| STARmap | Hundreds | submicrometric | 8 µm | 4 |
| Mass tomography | >50 | micrometric | Unlimited. Few mm in practice | |

[1] D. L. Coutu, K. D. Kokkaliaris, L. Kunz, T. Schroeder, Multicolor quantitative confocal imaging cytometry. *Nat Methods* **15**, 39-46 (2018)
[2] W. Li, R. N. Germain, M. Y. Gerner, Multiplex, quantitative cellular analysis in large tissue volumes with clearing-enhanced 3D microscopy (C. *Proc Natl Acad Sci U S A* **114**, E7321-E7330 (2017).
[3] Z. Zheng *et. al.*, A Complete Electron Microscopy Volume of the Brain of Adult Drosophila melanogaster. *Cell 174*, 730–743, July 26, 2018
[4] X. Wang *et al.*, Three-dimensional intact-tissue sequencing of single-cell transcriptional states. *Science* **361**, (2018).

### Supplementary Table 2

| Epithelial | | |
|---|---|---|
| Cytokeratin 14 (CK14) | polyclonal (Thermo Fisdher PA5) | Epithelial Marker |
| Pan-cytokeratin (panCK) | AE3 (Abcam, ab80826) | Epithelial Marker |
| Cytokeratin 19 (CK19) | Troma III (Hybridoma) | Epithelial Marker |
| Cytokeratin 8/18 (CK8/18) | C51 (Cell Signaling, 4546BF) | Epithelial Marker |
| HER2 | 29D8 (Cell Signaling 2165) | Epithelial Marker |
| Cytokeratin 5 (CK5) | EP1601Y (Abcam ab52635) | Epithelial Marker |
| E-Cadherin / P-Cadherin | 36/E-Cadherin (BD 610182) | Signal Transduction |
| Cytokeratin 7 (CK7) | RCK105 (BD 550507) | Signal Transduction |
| EGFR | D38B1 (Cell Signaling 4267BF) | Epithelial Marker |

| Chromatin | | |
|---|---|---|
| Histone H3 | D1H2 (Cell Signaling 4499BF) | Chromatin condensation |
| 3-Methyl-Histone3 | C36B11 (Cell Signaling 9733BF) | Chromatin condensation |
| Iridium | (Fluidigm) | DNA Binding |

| Other functions | | |
|---|---|---|
| Carbonic Anhydrase IX | polyclonal (R&D Systems AF2188) | Hypoxia |
| Cleaved Caspase3 | C92-605 (BD 559565) | Apoptosis |
| cleaved PARP | F21-852 (BD 552596) | Apoptosis |
| Ki-67 | B56 (BD 556003) | Proliferation |

| Phenotyping of other cells | | |
|---|---|---|
| Vimentin | D21H3 (Cell Signaling 5741BF) | Cytoskeleton. Mesenchymal. |
| HLA-ABC | W6/32 (iolegend 311402) | Antigen Presentation |
| Alpha-SMA | 1A4 (Abcam ab7817) | Smooth Muscle, Perycytes |
| Fibronectin | 10/Fibronectin (BD 610078) | Extracellular |
| Syndecan-1 | MI15 (Biolegend 356502) | Extracellular - Membrane |
| Collagen I | polyclonal (Abcam ab34710) | Extracellular |

| Immune Cell Phenotyping | | |
|---|---|---|
| CD20 | L26 (E-Bioscience \| 14-0202-82) | B Cells |
| CD44 | IM7 (BD 550538) | Myeloid, mesenchymal |
| CD45 | 2B11 (eBioscience 14-9457-82) | Immune Cells |
| CD68 | KP1 (E-Bioscience 14-0688-82) | Monocyte/Macrophage |
| CD8 alpha | C8/144B (E-Bioscience 14-0085-82) | Cytotoxic T Cells |
| Myeloperoxidase MPO | polyclonal (Dako A0398) | Granulocytes |
| CD3 epsilon | D7A6E (Cell Signaling 85061) | T Cells |

| Signal Transduction | | |
|---|---|---|
| pS6 | D57.2.2E (Cell Signaling 4858) | Signal Transduction |
| pErk1/2 | 20A (BD 612359) | Signal Transduction |
| phospho-H3 | HTA28 (Biolegend 641007) | Signal Transduction |

Supplementary Table 3

| | Multi-parametric | Cell Data analysis | mcd file format | License | Image registration | Intensive 3D rendering |
|---|---|---|---|---|---|---|
| histoCAT+3D | Yes | Yes | Yes | A-GPL | Yes, adapted to IMC | Yes |
| Imaris | Limited | Yes (extra paid module) | No | Commercial | Yes | Yes |
| CellProfiler | Yes | Yes | No | BSD-3 | No | ~ (only 3.0 and memory limited) |
| Ilastik | Yes | Yes | No | GPL | No | Yes |
| Fiji | Yes | Limited (plug-ins) | Yes (mcdtools†) | OSS | Yes | ~ (plug-in dependent) |

95ºC Antigen Retrieval

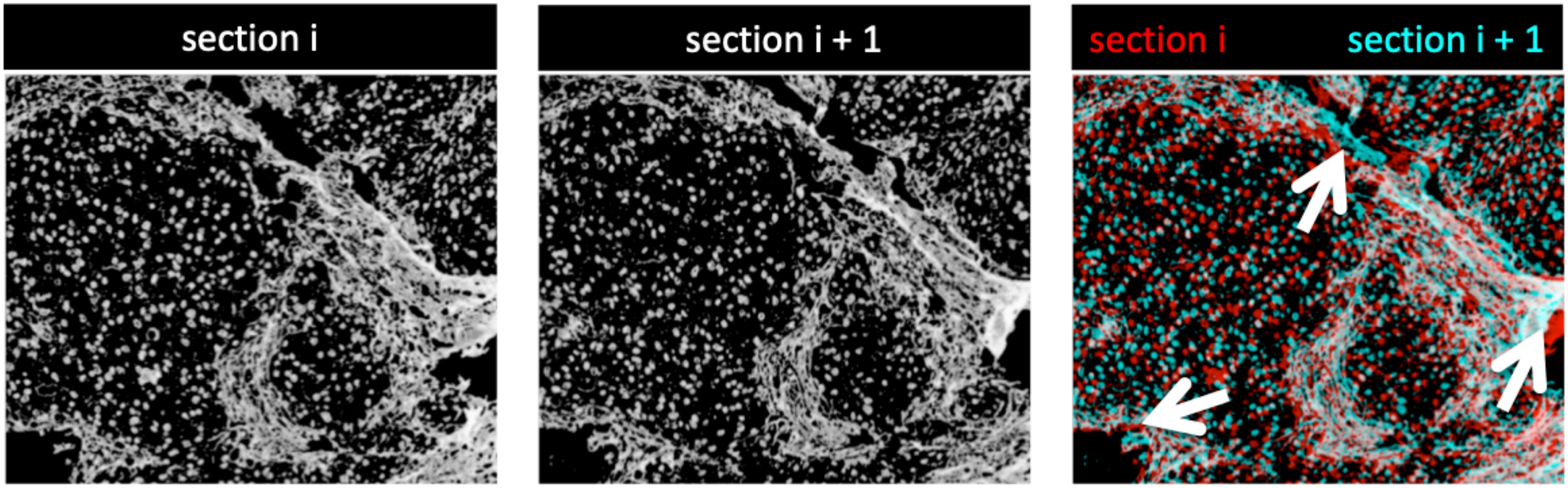


80ºC Antigen Retrieval

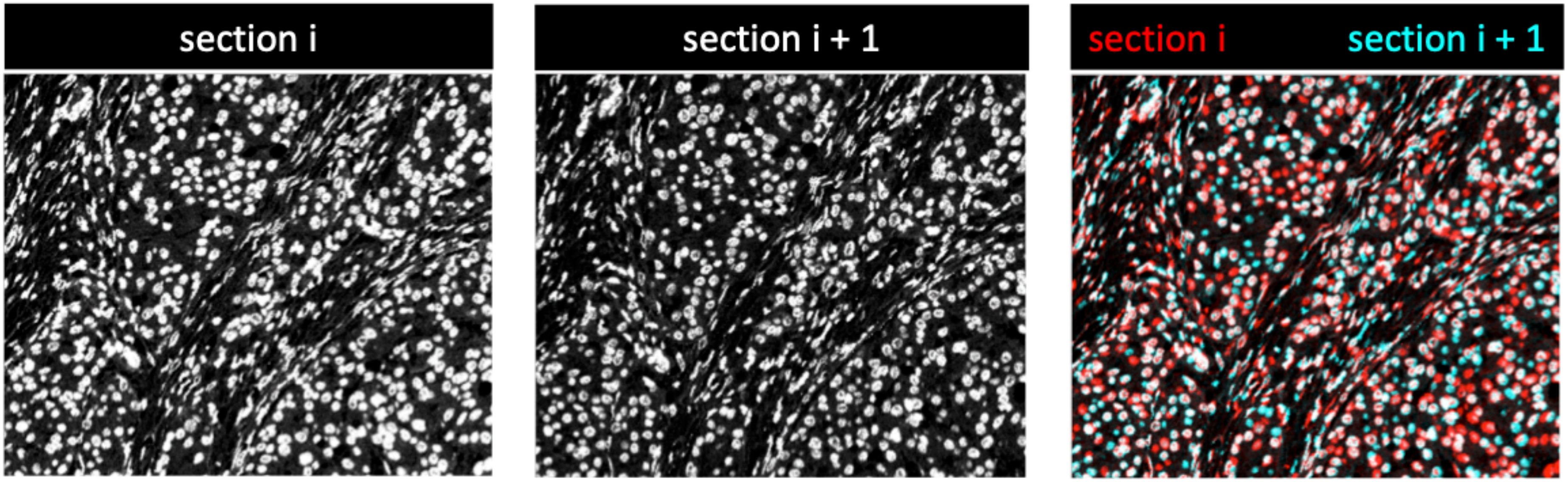

Supplementary Figure 2

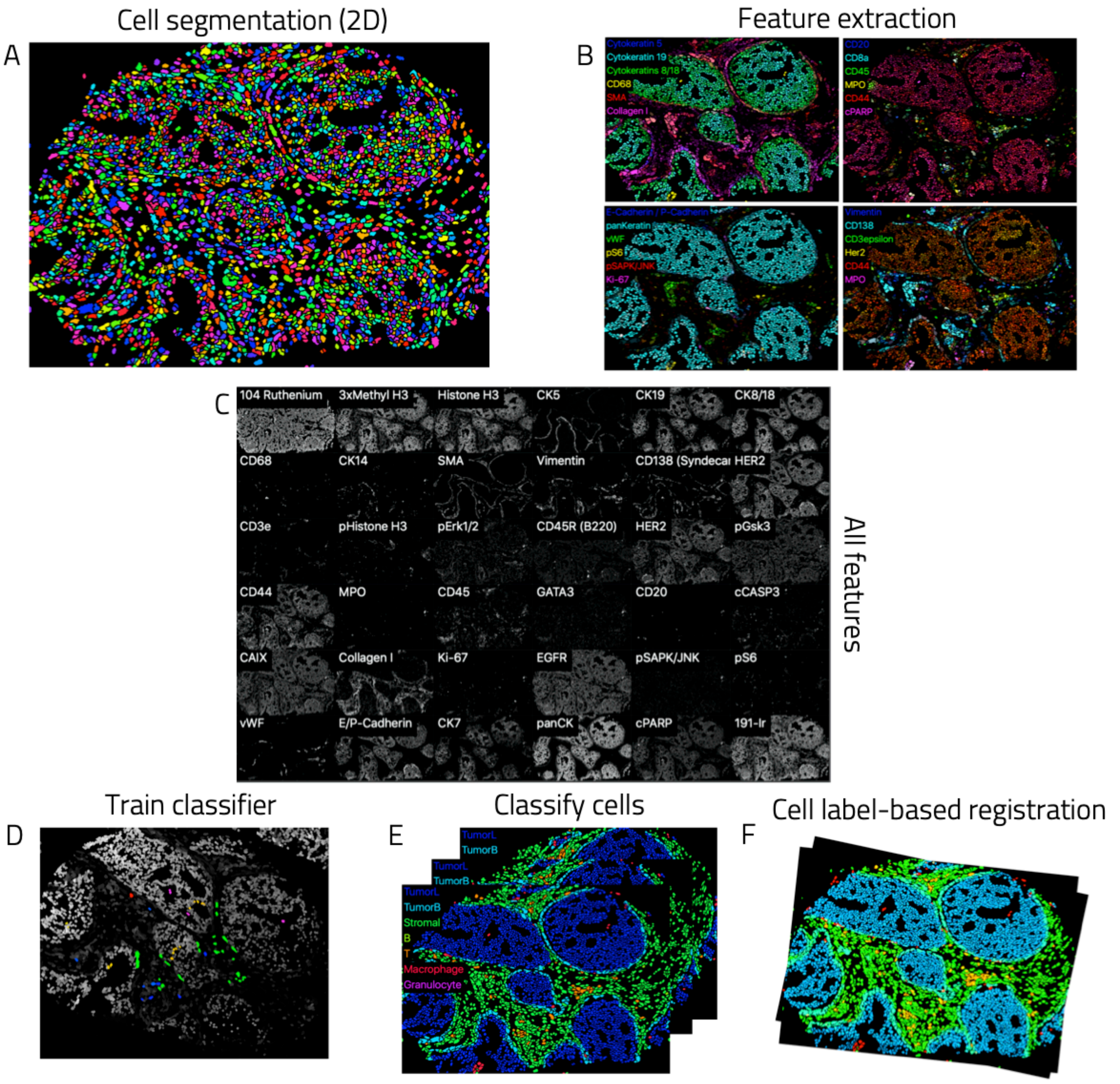

Supplementary Figure 3

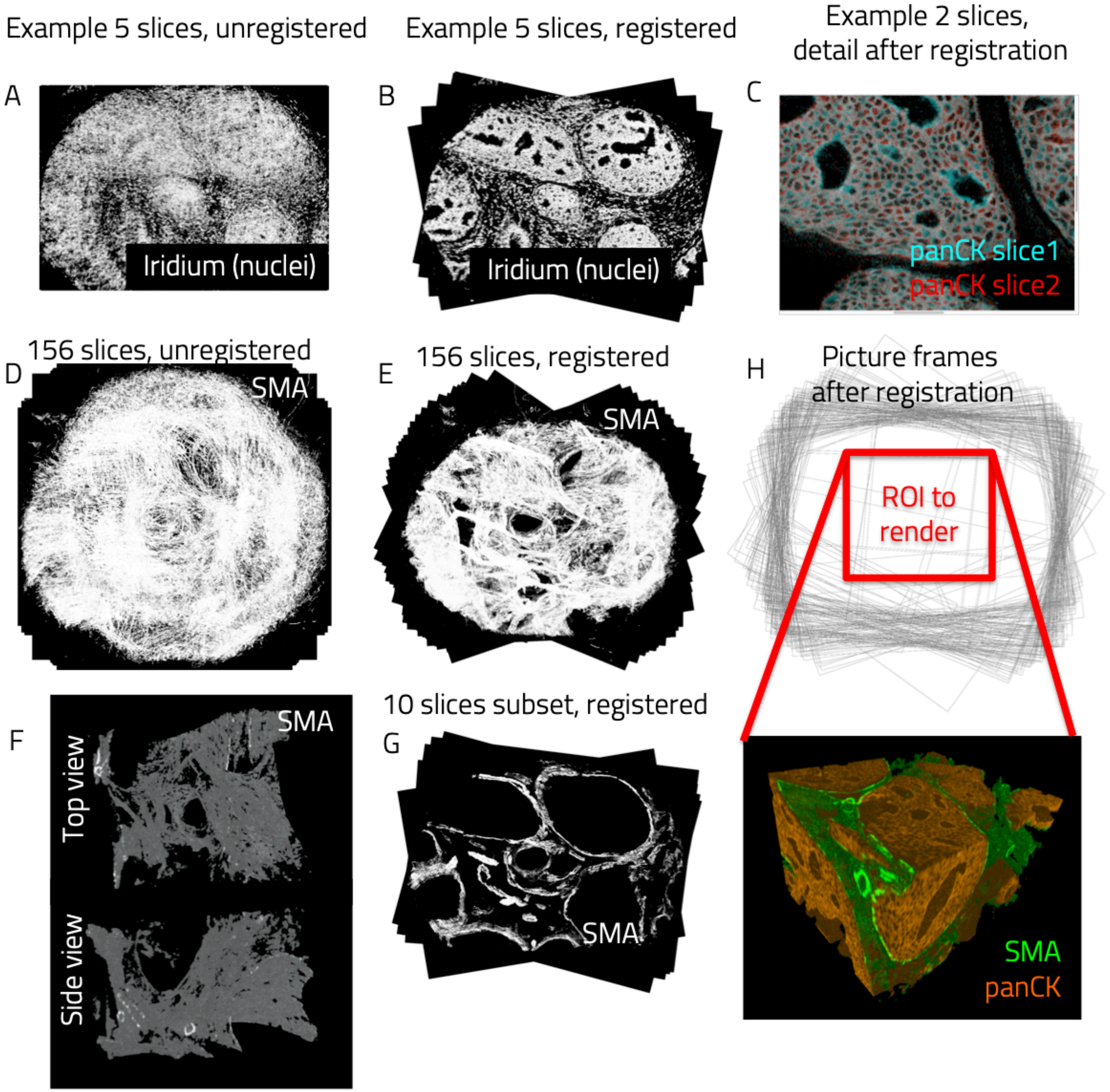

Supplementary Figure 4

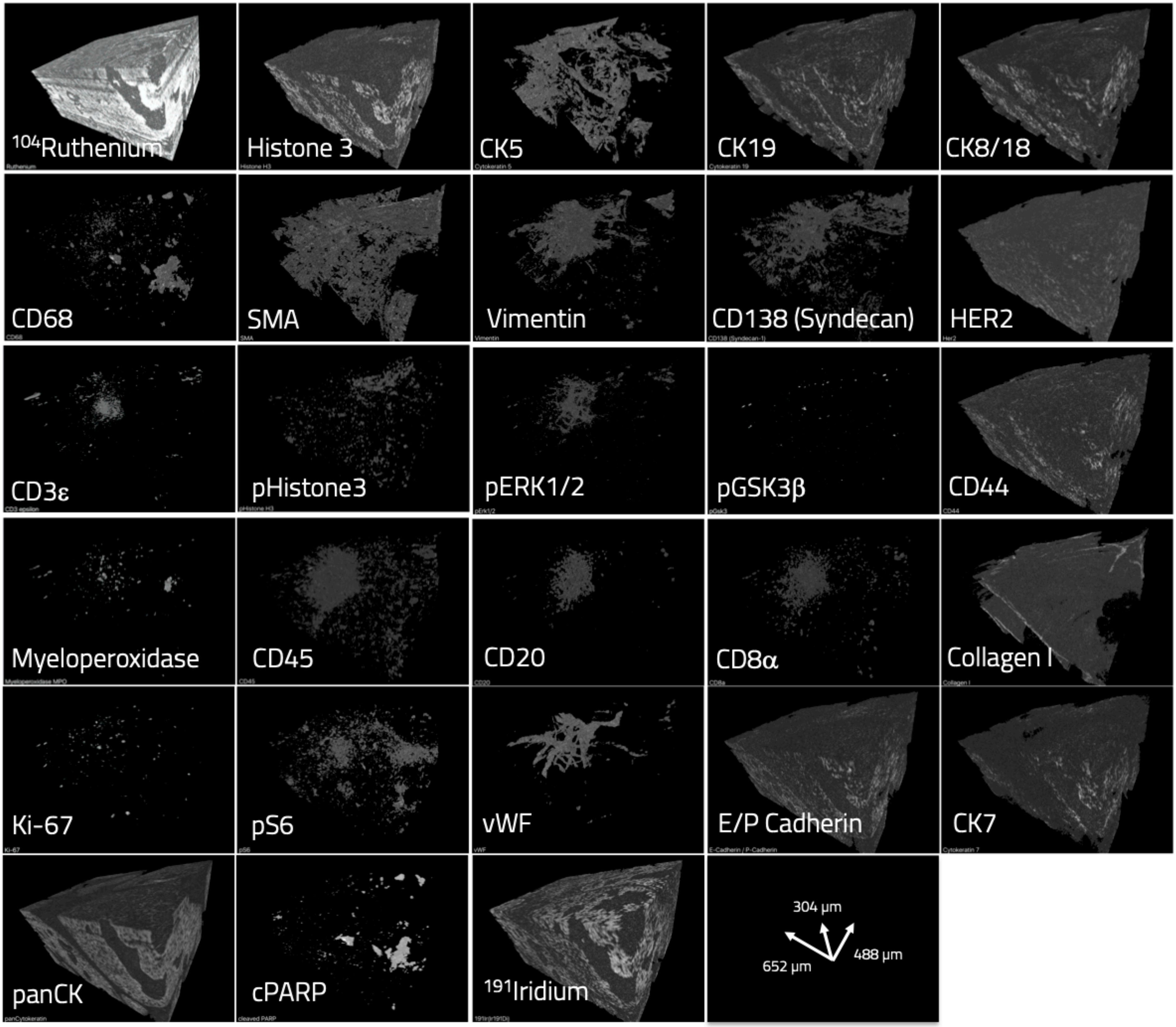

Supplementary Figure 5

No filter

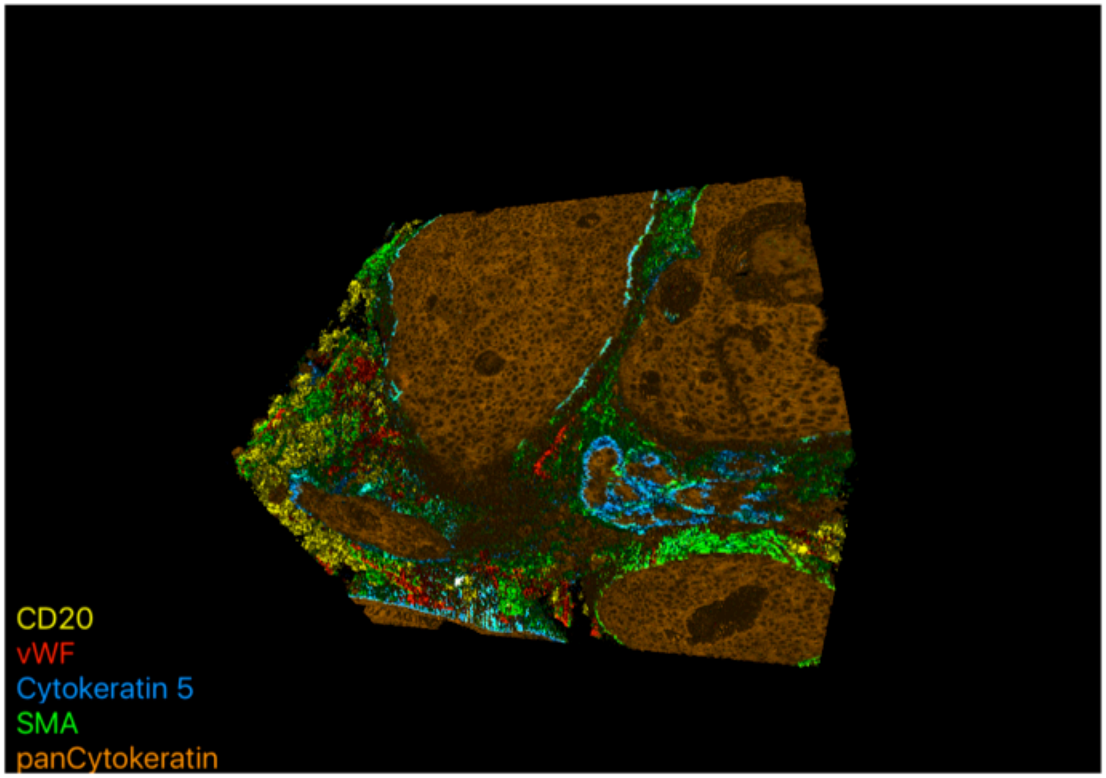


Gaussian filter

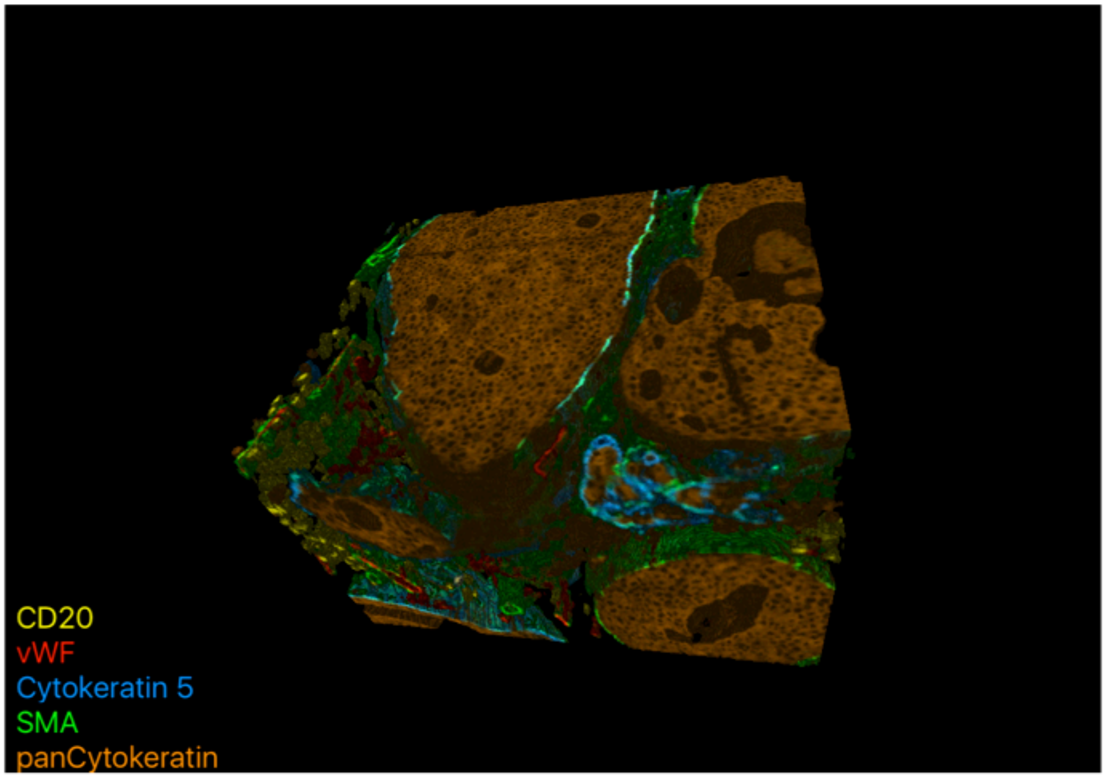

Supplementary Figure 6

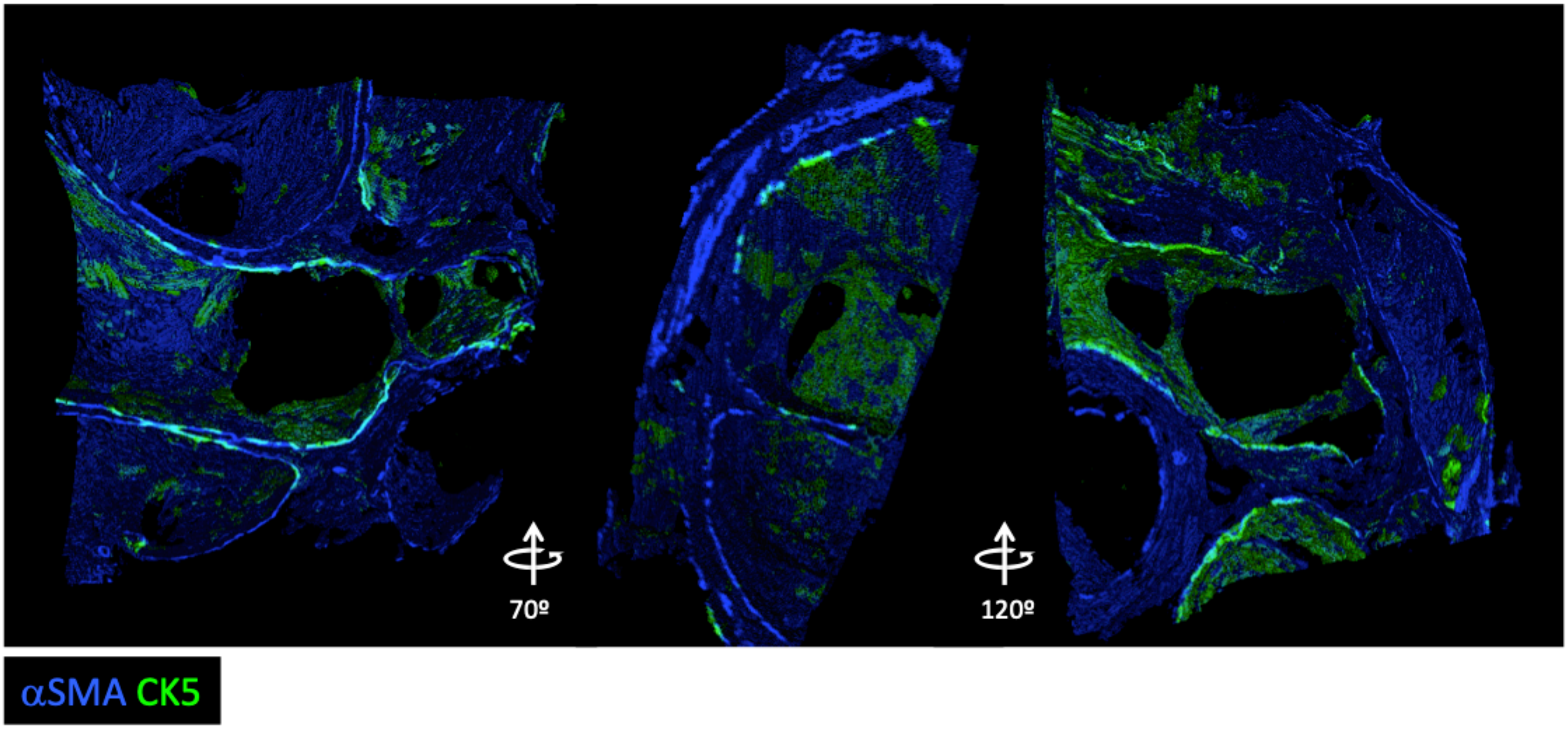

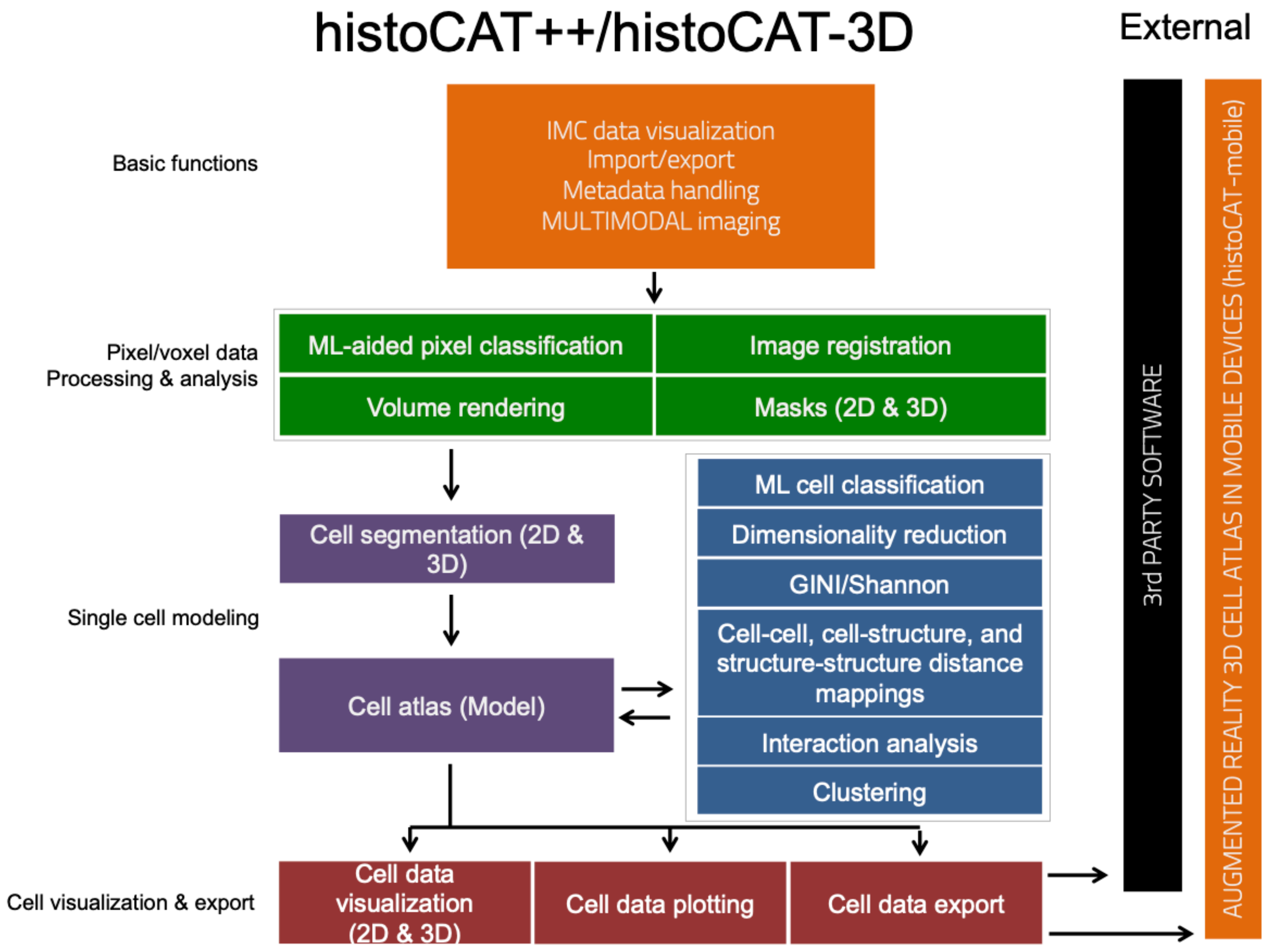
histoCAT++/histoCAT-3D
External
Basic functions
IMC data visualization
Import/export
Metadata handling
MULTIMODAL imaging
Pixel/voxel data
Processing & analysis
ML-aided pixel classification
Image registration
Volume rendering
Masks (2D & 3D)
Single cell modeling
Cell segmentation (2D & 3D)
Cell atlas (Model)
ML cell classification
Dimensionality reduction
GINI/Shannon
Cell-cell, cell-structure, and structure-structure distance mappings
Interaction analysis
Clustering
Cell visualization & export
Cell data visualization (2D & 3D)
Cell data plotting
Cell data export
3rd PARTY SOFTWARE
AUGMENTED REALITY 3D CELL ATLAS IN MOBILE DEVICES (histoCAT-mobile)

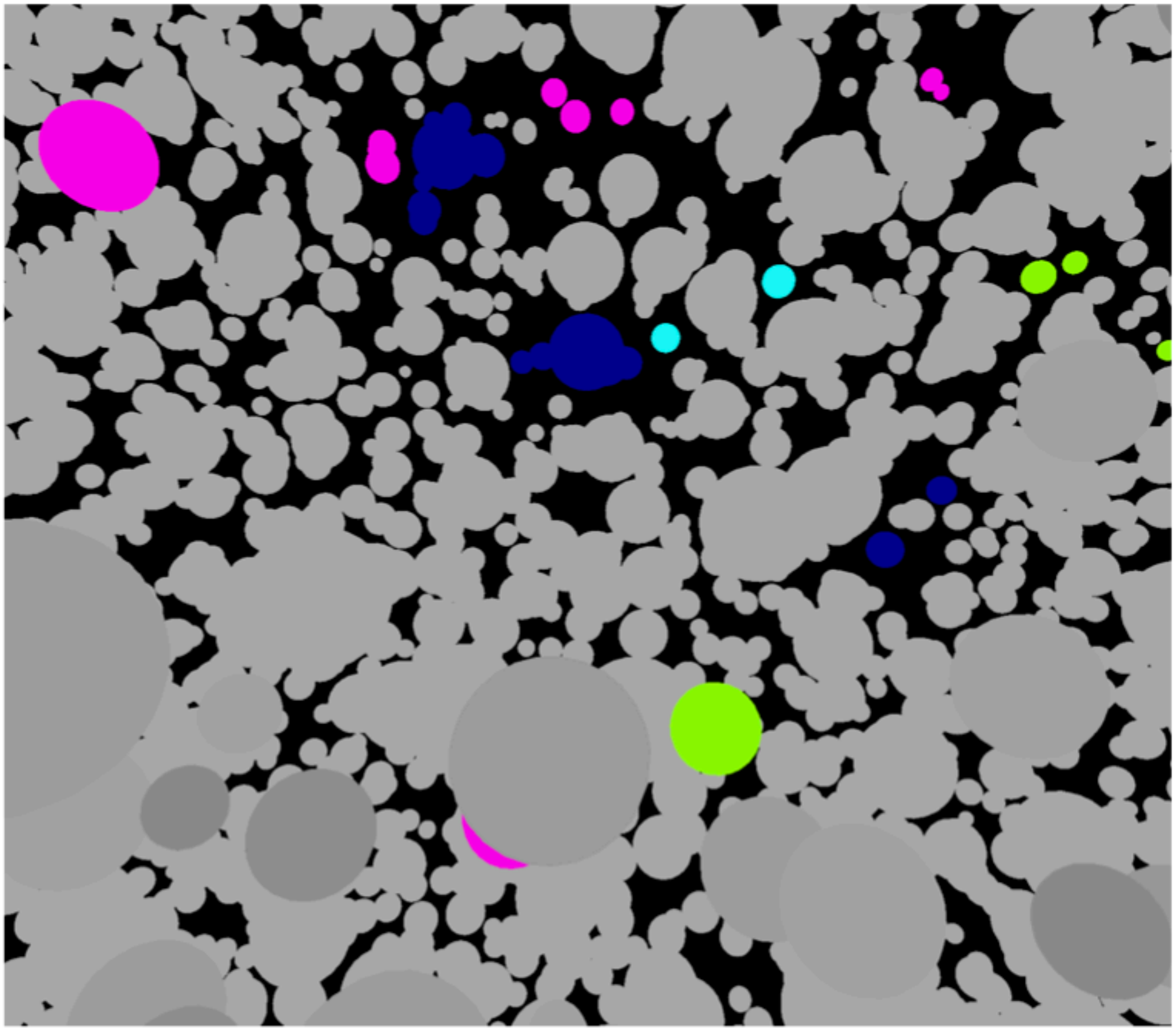

Supplementary Figure 9

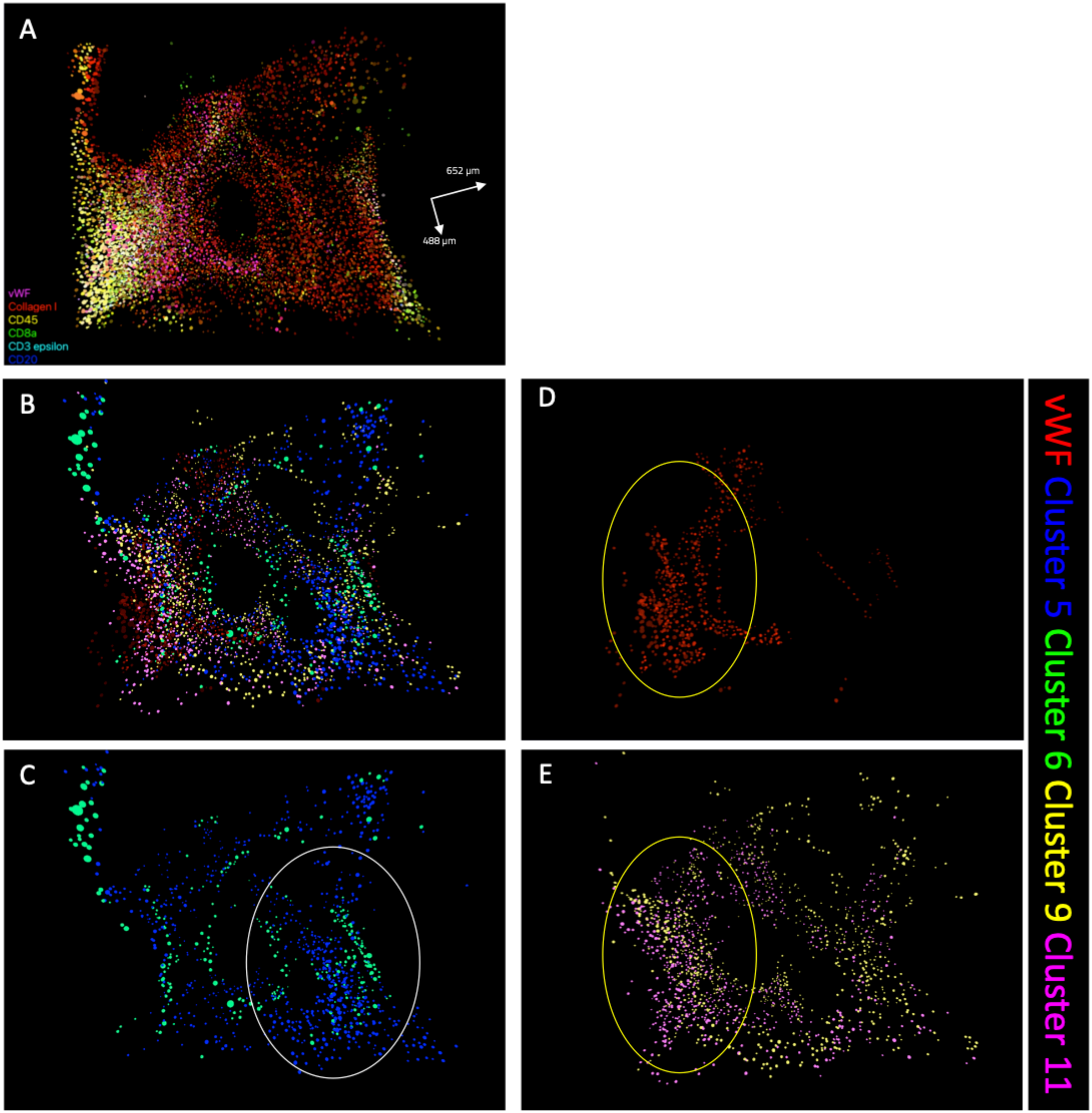

Supplementary Figure 10

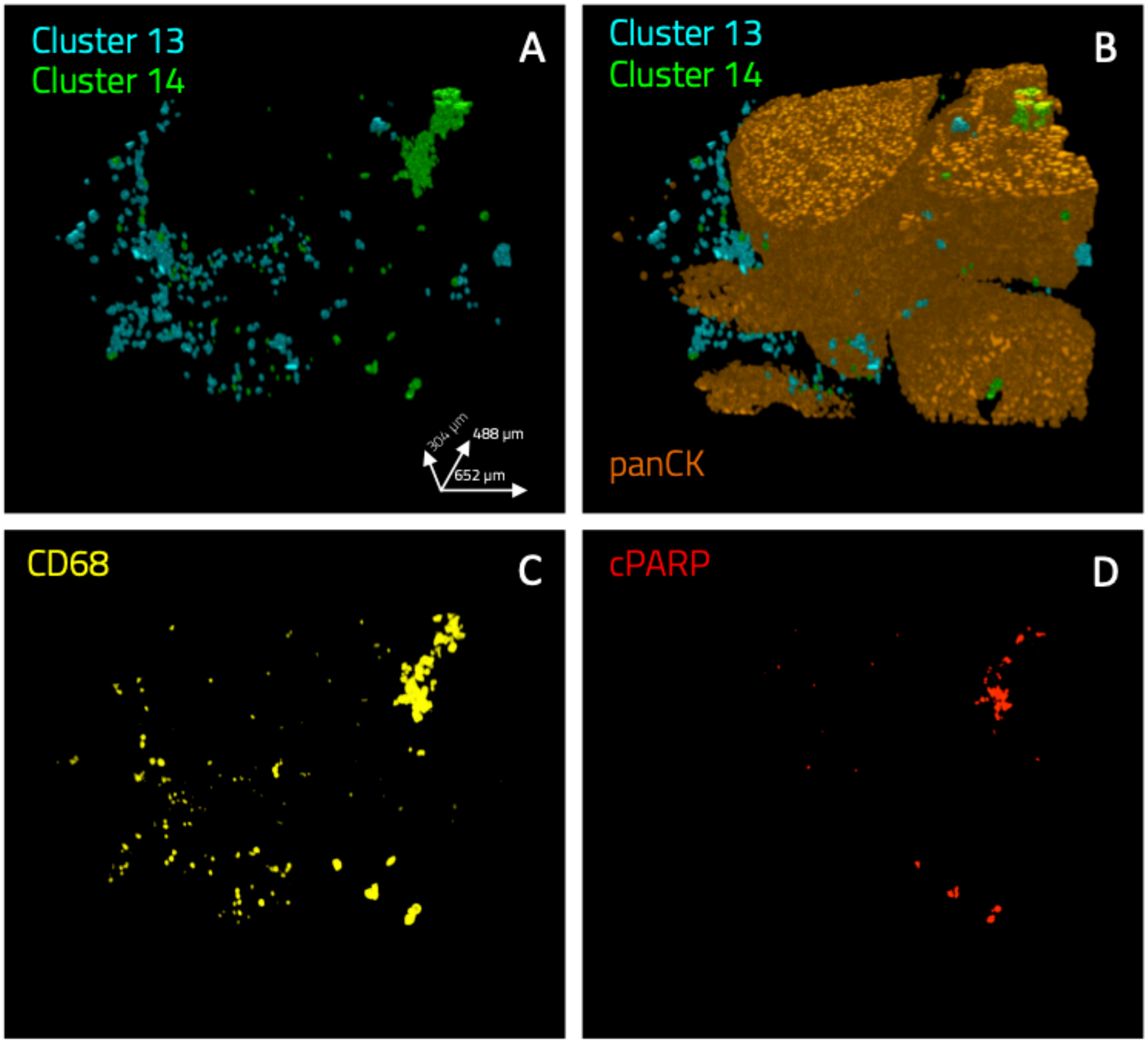

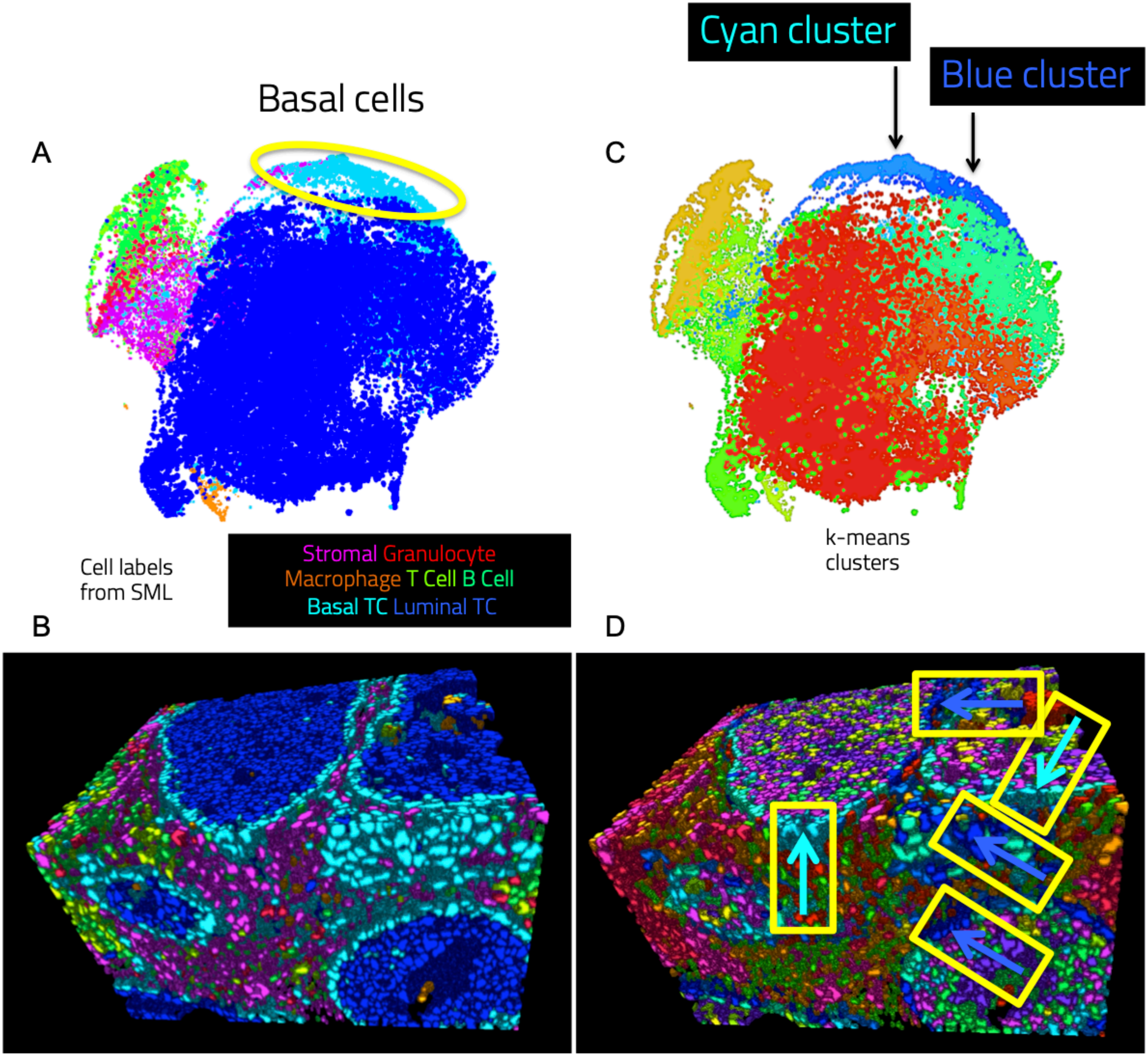
Basal cells
A
Cyan cluster
Blue cluster
C
Cell labels
from SML
Stromal Granulocyte
Macrophage T Cell B Cell
Basal TC Luminal TC
k-means
clusters
B
D

Supplementary Figure 12

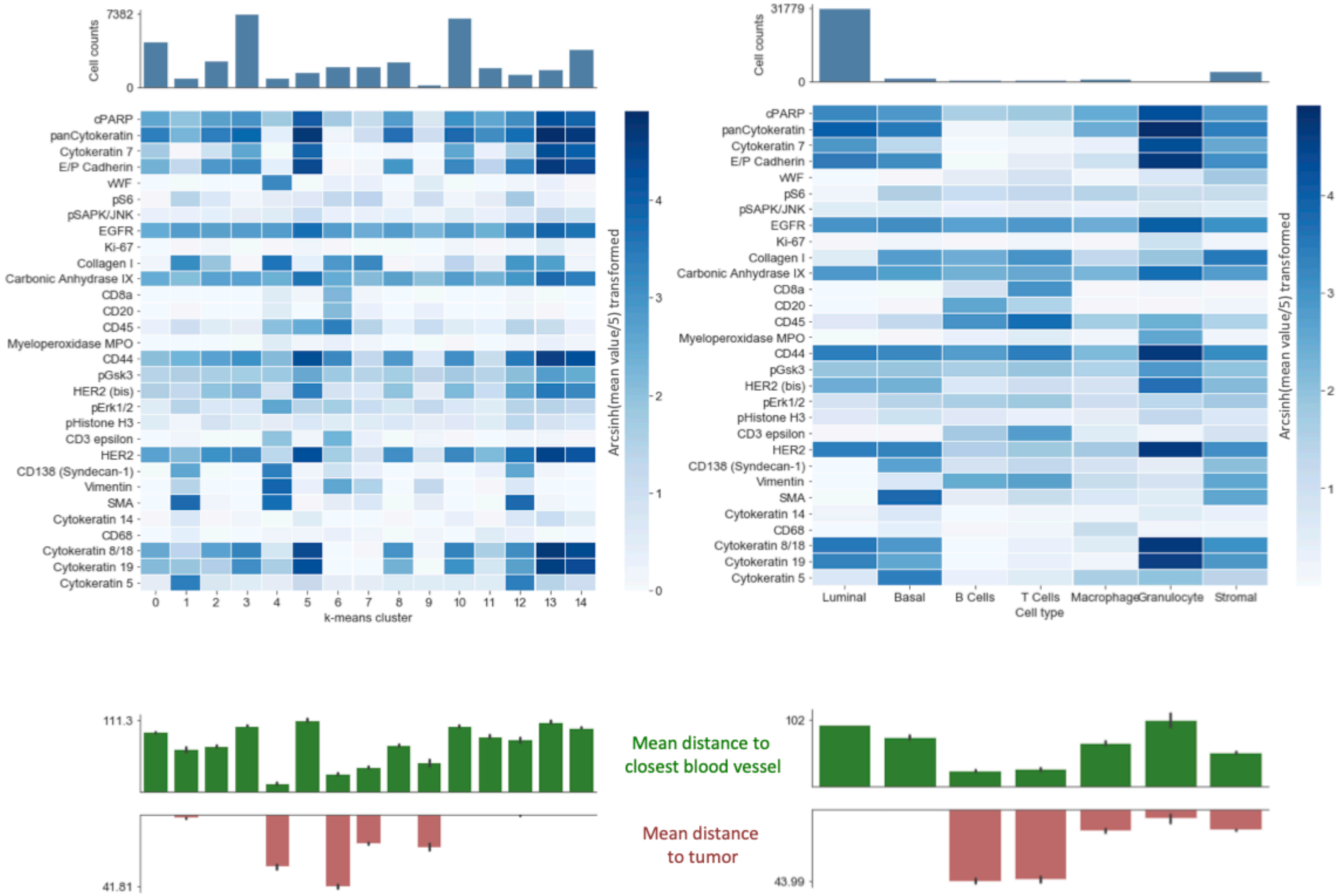

### Supplementary Figure 13

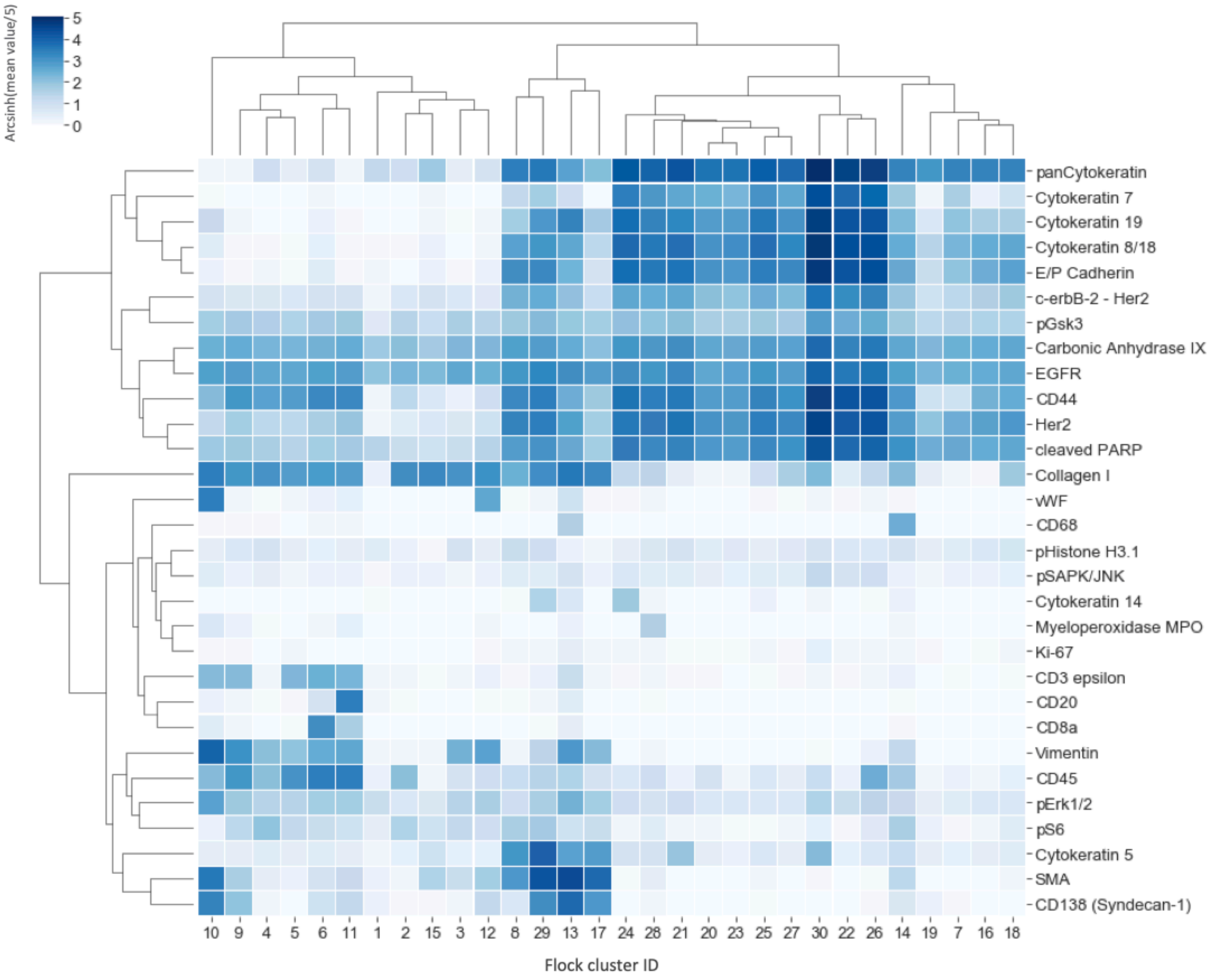

### Supplementary Figure 14

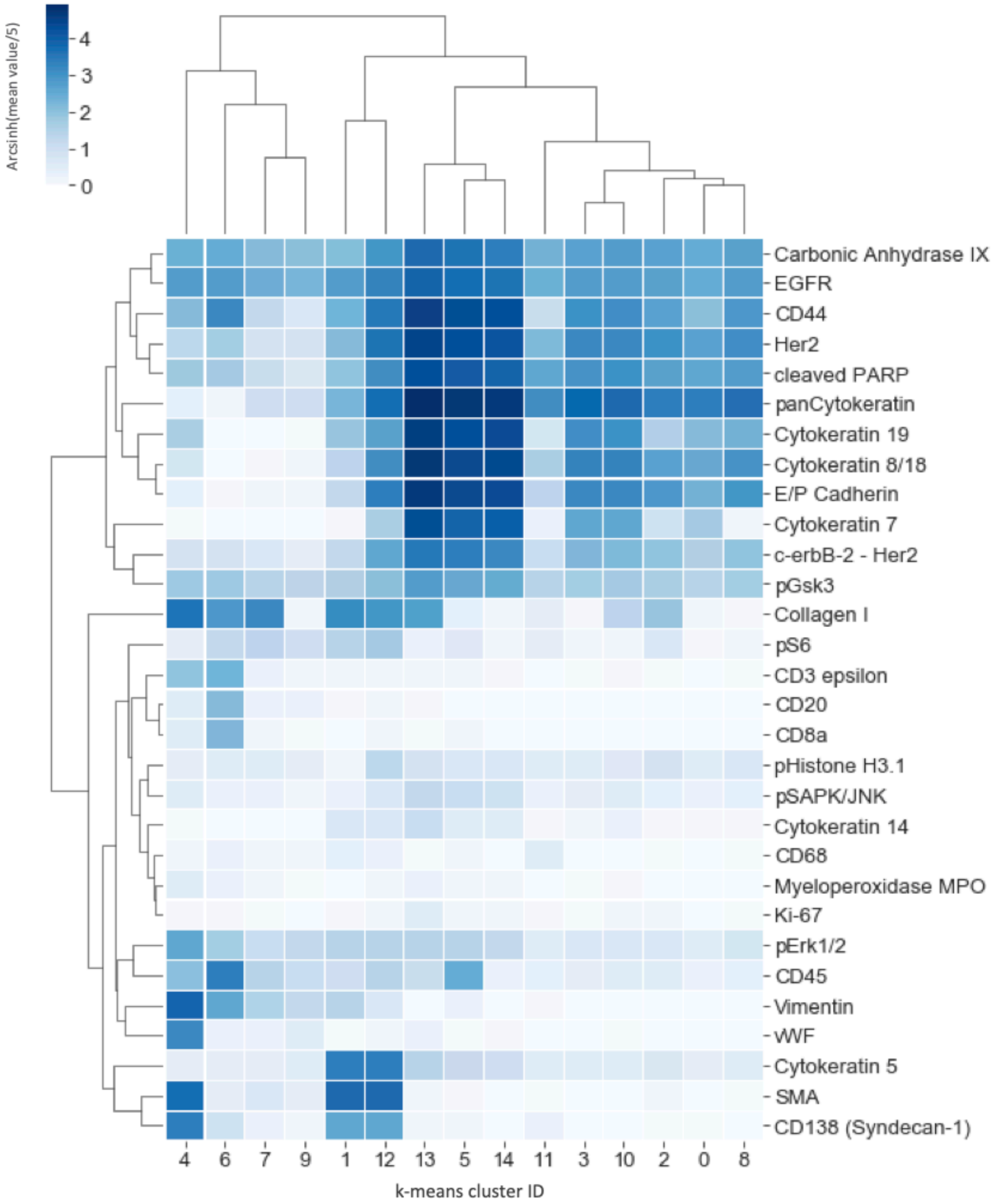

Supplementary Figure 15

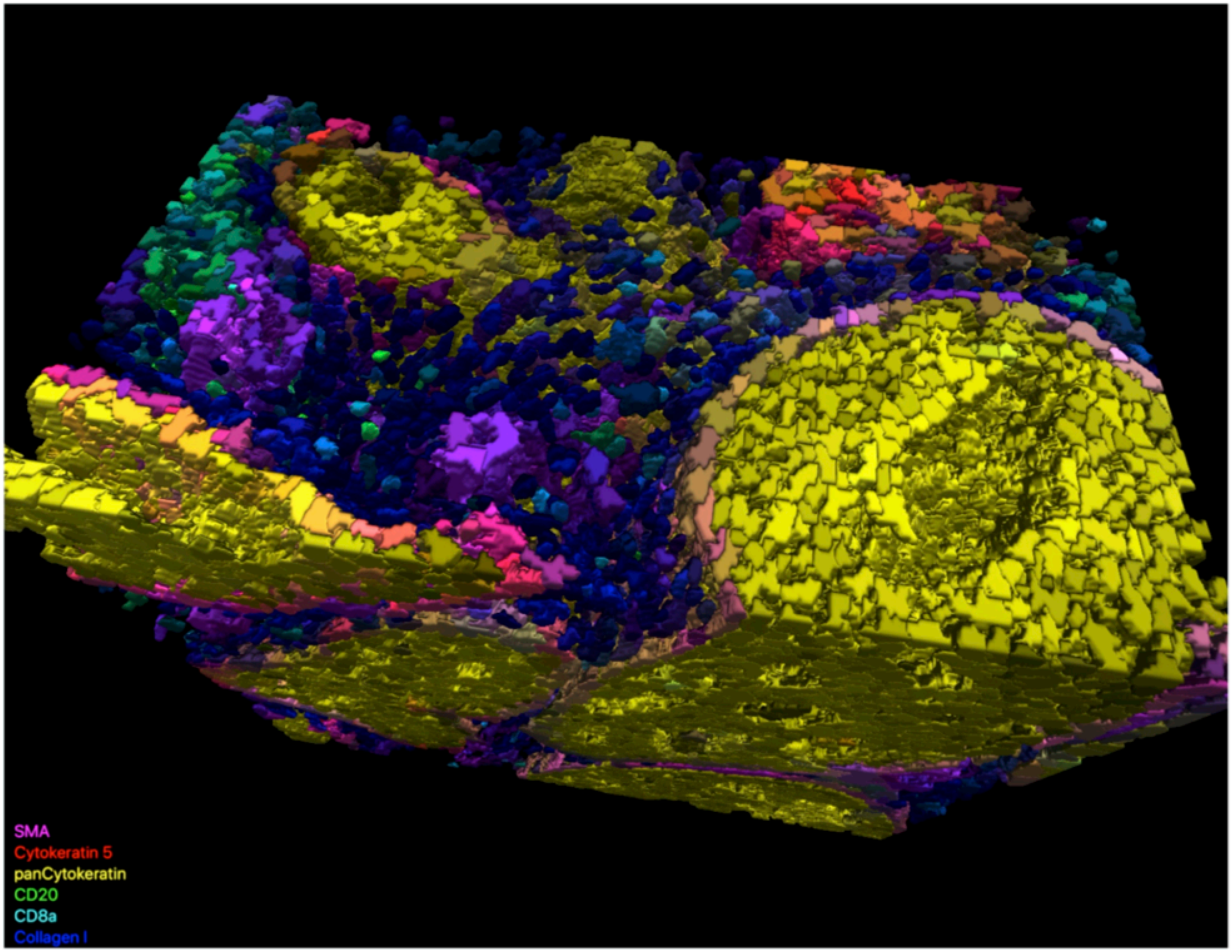
