## Supplementary Methods for "Highly multiplexed molecular and cellular mapping of breast cancer tissue in three dimensions using mass tomography"

Breast cancer tissue

We use archived material form the tissue biobank at the University Hospital, Zürich. Tissues of interest were screened using traditional Hematoxylin and Eosin stained sections from Formalin-Fixed, Parafin-Embeded (FFPE) tissue blocks. Once regions of interest were identified by the pathologist, a 1mm diameter cylindrical punch was obtained in a similar fashion to that of tissue microarray elaboration. One sample was used for this study. This sample was part of a study which was approved by the ethics committee of the Canton Zurich (PB_2016-00811).

Antibody Testing and Validation

A panel of antibodies was designed using AirLab (Catena et. al, Genome Biology, 2016). All antibodies in the panel (Supplementary Table 1). were conjugated to metals using the MaxPar X8 Multimetal Labeling Kit (Fluidigm) according to the manufacturer's instructions. Antibodies were initially tested and titrated for IMC in single breast cancer FFPE sections. Antibodies displaying expression patterns consistent with the literature and a sufficient signal intensity, were utilized. Finally, the entire antibody panel was used to stain consecutive batches of 2 micra slices obtained by ultramicrotomy (as described below) to generate 3D reconstructions.

Mass Tomography Protocol

Sample sectioning (1 day)

1. Obtain samples in paraffin. There are two options, both starting from paraffin blocks (see how to embed samples in paraffin elsewhere)
   - Cut prism of about 1 x 1 mm cross-section and between 5 to 10 mm height.
   - Use cylinder for Tissue Microarray production to get a cylinder punch of around 1mm diameter and with the block thickness.

2. Re-cast tissue into EPON-silicon molds.
   2.1. Dump prism/cylinder into molten paraffin container and clear remaining paraffin
   2.2. Place tissue into silicon molds (normally used for EPON embedding) with the cylinder axis along the axis of the mold. Fill mold with molten paraffin in excess and quickly move to cold surface to solidify paraffin.
   2.3. Wait for 20/30 minutes at 4C
   2.4. Remove block from mold and with blade carefully remove salient paraffin borders.

3. Mount block in ultramicrotome capsule holder

4. Use a diamond trimming knife to trim a trapezium shape with tissue exposed in the salient block face
5. Change to Histo-Jumbo Diamond knife (Diatome) with plastic-sealed hole in bottom. Insert needle connected to ultrapure water syringe through the bottom whole. Fill diamond knife boat with ultrapure water. Adjust level to edge of knife carefully using the syringe embolus.
6. Carefully approach knife to sample. Setting the step length to 2 micra, trim again the block face sections are cut through the whole block face and the whole tissue of interest is exposed. This is necessary as angles change a little between trimming with the glass knife and changing to the HistoJumbo diamond knife.
7. Clean blade border with Styrofoam rods. Introduce Superfrost histological glass slides in boat of Jumbo knife. Adjust water levels again if necessary.
8. Carefully cut ribbons, with eyelash pole, aggregate/rack ribbons, approach them to opposite side of knife.
9. Once ready to collect, with one hand slowly empty water from boat. With the other, use the eyelash pole to direct ribbons to central area of the glass slide.
10. Carefully collect slide out of the knife's boat, tap gently to remove remaining water.
11. Place slide on warm metal plate (30C) for 2 hours to tether tissues to the glass slide.

Tissue preparation and antigen retrieval (~5h)

1. Place slides in clean xylene. Incubate for at least 2h
2. Change slides to a cuvette with fresh xylene. Incubate at least 1h.
3. Changes slides to a Xylene:Ethanol (1:1) koplin jar. Incubate for 10 minutes.
4. Move slides to a Ethanol 100% koplin jar. Incubate for 10 minutes.
5. Hydrate tissue by moving tissues through Ethanol 96%, 90%, 80%, 70% to ultrapure water. 5 minutes each.
6. From water, move slides to cuvette with Antigen retrieval buffer (10mM Tris-HCl buffer, pH 10.0, also know as HIER buffer)
7. Perform antigen retrieval in pressure cook (Medite) at 80ºC for 80 minutes.
8. Cool down slides slowly (snap temperature changes can cause deformation of sections)
9. Move slides to water quickly, then to TBS or PBS. Incubate for 5 minutes at RT

Sample staining (~20h)

1. Paint around tissue the stainable area with hydrophobic pen.
2. Add blocking buffer (PBS or TBS with 1% Horse Serum and 1% FcBlock)
3. Incubate for 30 minutes at RT
4. Add antibody mix (diluted in blocking buffer). Incubate overnight at 4C. You may need to incubate in more than one round if antibody amounts exceed 50% of the staining volume. For instance, for 200 µL staining volume, if the antibody volumes add up to 140 µL, split the panel in 2, prepare half the panel

(presumably ~70 µL) and add blocking buffer to 200 µL, incubate overnight, and then do the same with the other half of the panel. It is not necessary to wash the tissues in between antibody rounds.
5. Wash 3x in TBS or PBS for 5 minutes each wash.
6. Add Iridium intercalator, incubate for 2 to 5 minutes.
7. Wash iridium intercalator in PBS for 10 seconds. Do not use TBS for this wash.
8. Add Ruthenium tetroxide 1:500 / 1:1000 (Starting from a 0.4% commercial solution). Dilute the Ruthenium tetroxide only in PBS (TBS and serum components react).
9. Incubate for 3 to 5 minutes.
10. Wash Ruthenium tetroxide in ultrapure water directly.
11. Air-dry (be careful not to be aggressive with the airstream over the tissue as you could get deformation).

Acquisition (3-5 days, depending on number of sections and size of area obtained)

1. Data acquisition was performed on a Helios time-of-flight mass cytometer (CyTOF) coupled to a Hyperion Imaging System (Fluidigm). Prior to laser ablation, optical images of slides were acquired using the Hyperion software, and the areas of all consecutive tissue sections to perform acquisition were selected as described above. Selected areas for ablation were bigger than the actual area of interest to account for loss of overlapping areas amongst sections due to cumulative rotations (Supplementary Fig. 3).
2. Laser ablation was performed at a resolution of approximately 1 µm with a frequency of 200 Hz. To ensure performance stability, the machine was calibrated daily with a tuning slide spiked with five metal elements (Fluidigm).
3. All data was collected using the Fluidigm CyTOF software v6.7. Raw data is available at http://www.bodenmillerlab.org/catena_et_al_mass_tomography/index.php

Reconstruction and analysis (1-2 days)

The software package histoCAT++/3D, presented in this manuscript was used to perform all the preprocessing and analytical steps. A detailed manual for histoCAT++/3D is available at www.bodenmillerlab.org/histoCAT_manual. Code can be found at github.com/BodenmillerGroup/histoCAT3D/blob/master/3DIMC.

The following steps were followed with histoCAT++/3D (Supplementary figures 2-3 and all main figures):

1. Channel compensation, using the compensation matrix already published by our group (Chevrier et. al, Cell Systems 2018). The code implementing channel compensation in histoCAT++/3D can be found in the file IMCImageStack.m,
2. Channel equalization across the full stack of 168 images. Implemented methods can be found in the file IMCImageStack.m.

3. Reassembly of images corresponding to sections not recorded in a single acquisition. The 156 slices analyzed correspond to 168 IMC acquisitions, with 10 cases were a section is contained in 2, or 3 different IMC images. See files IMC3DMask.m and IMC3DMaskComputations.m
4. Nuclear and cytoplasm segmentation (using iridium and pan-cytokeratin signals) using a watershed algorithm. (IMCWatershedSegmenter.m, method extractMaskFromRender:channels:dictChannel:framingChannel:dictSChannel :threshold:gradient:minKernel:expansion:name:) This method is a custom implementation of a watershed algorithm that takes an image, a set of channels (that are summated, normally only nuclear channels are included here), a so-called framing-channel (substrated from the previous summation. This helps separating cells that are too close or touching, for this channel normally a membrane or cytoplasmic channel such as HER2 or pan-Cytokeratin is used), a threshold value for the watershed (0 to 1, typically 0.1, to set the percentage of intensity for which the algorithm will stop), gradient (0.02-0.1 is the watershed intensity threshold decrease for every step of the basins filling), minimum kernel size for a minimum number of connected pixels to consider a cell, and expansion (1, 2 normally, for extending the radius, in pixels, of the resulting mask once the algorithm is done). During the algorithm, all blobs acquiring the minKernel area, are assigned a unique ID. The resulting segmentation mask contains 0 where there is no cell, and the ID of each of the segmented cells.
5. Molecular and morphological feature extraction for all cells in 2D. (IMCComputationOnMask.m, method extractDataForMaskOperation:computations processedData:). This method computes mean, median, and standard deviation for all the channels in the original image files within each of the segmented cells.
6. Training of a random-forest classifier for 7 different cell populations (Luminal epithelial, basal epithelial, B cells, T cells, macrophages, granulocytes, and other stromal cells) by clicking cells with the corresponding antigen expression in three different images. (IMCCellTrainer.m, IMCCellTrainerTool.m, IMCRandomForests.mm, RandomForests.cpp). User can define labels and click over cells to train a classifier (build on top of OpenCV library´s implementation) which is then used to classify all cells in the datasets (see manual for screenshots).
7. Classifying all cells in 157 assembled sections into the 7 categories
8. Perform registration of all images with a cell-label matching algorithm. This algorithm iterates over different rotations and translations of the moving image and computes a distance versus the target image using simple label matching as a metric instead of distance metrics over the intensities. (See IMCRegistration.m)
9. Rasterize of the signal of all realigned images and collation of all stack data into a voxel model.
10. Render of the voxel model with different channels and colors. Using the Metal2 Apple´s framework (IMCMetal**Renderer.m files)
11. 3D segmentation using a custom 3D watershed algorithm (IMC3DMask.m).

12. Segmentation of tumor and blood vessel structures using a channel threshold filter and the aforementioned modified 3D-watershed algorithm (IMC3DMask.m).
13. Molecular and morphological feature extraction for all cells in 3D. In an analogous way to what described in point 5 above for 2D. (IMC3DMask.m)
14. tSNE dimensionality reduction. (Reimplemented from van der Maaten, https://lvdmaaten.github.io/tsne/ and wrapped for Objective-C, see files IMCBhSNEOperation.mm, bhsne.cpp, IMCCellBasicAlgorithms.m)

FLOCK and k-means clustering. (IMCFlockOperation.m, flock.c). Reimplemented from the C version (https://sourceforge.net/projects/immportflock/files/FLOCK_version_1/) published by Richard H. Scheuermann et al. (Journal of Immunology, 2009)

15. Interaction matrix calculation.
16. Distance to vessel calculation for all cells. (IMC3DMask.m) Simply calculates the number of cell-cell interactions amongst cells by type (using cell labels generated as described in points 6-7), the number of expected interactions according to the cell type abundances, and finally, the ration of observed versus expected interactions.
17. Distance to tumor calculation for all cells. (IMCCombineMasks.mm)
18. 3D cell data rendering as voxels. (IMCMetalViewAndRenderer.m)
19. 3D cell data rendering as spheres. (IMCMetalSphereRenderer.m)
20. 3D cell data rendering as striped-spheres. (IMCMetalSphereStripedRenderer.m)
21. 3D cell data rendering as isosurfaces. (IMCMetalPolygonizedRenderer.m) after creating a mesh using the marching cubes algorithm (IMC3DMask.m)
22. Video generation. (IMCVideoCreator.m)
23. Exporting cell data for histoCAT mobile (https://github.com/BodenmillerGroup/histoCATmobile) and for further analysis using python scripts.

All data, the histoCAT++/3D workspace, and analysis files are available at http://www.bodenmillerlab.org/catena_et_al_mass_tomography/index.php

The histoCAT-3D software package is available at: http://www.bodenmillerlab.com/research-2/histocat-3/.
